# A-SOiD, an active learning platform for expert-guided, data efficient discovery of behavior

**DOI:** 10.1101/2022.11.04.515138

**Authors:** Jens F. Tillmann, Alexander I. Hsu, Martin K. Schwarz, Eric A. Yttri

**Author notes:** These authors contributed equally (Jens F. Tillmann, Alexander I. Hsu). These authors contributed equally (Martin K. Schwarz, Eric A. Yttri).

## Abstract

To identify and extract naturalistic behavior, two schools of methods have become popular: supervised and unsupervised. Each approach carries its own strengths and weaknesses, which the user must weigh in on their decision. Here, a new active learning platform, A-SOiD, blends these strengths and, in doing so, overcomes several of their inherent drawbacks. A-SOiD iteratively learns user-defined groups and can considerably reduce the necessary training data while attaining expansive classification through directed unsupervised classification. In socially-interacting mice, A-SOiD outperformed other methods and required 85% less training data than was available. Additionally, it isolated two additional ethologically-distinct mouse interactions via unsupervised classification. Similar performance and efficiency were observed using non-human primate 3D pose data. In both cases, the transparency in A-SOiD’s cluster definitions revealed the defining features of the supervised classification through a game-theoretic approach. Lastly, we show the potential of A-SOiD to segment a large and rich variety of human social and single-person behaviors with 3D position keypoints. To facilitate use, A-SOiD comes as an intuitive, open-source interface for efficient segmentation of user-defined behaviors and discovered sub-actions.

## Introduction

Naturalistic behaviors, particularly social interactions, provide a rich substrate to understand the brain and its decisions. Cutting-edge machine learning algorithms now enable researchers to capture the movement of individual body parts with markerless pose estimation [1–7]. These pose estimation algorithms can then be readily used to extract behavioral expressions in a previously unmatched level of detail and temporal resolution [5, 6, 8–15].

Pose estimation data can be used to train an algorithm to reproduce expert human annotation in an automated fashion. A major advantage of these supervised approaches is the direct control over the initial definition of the behavioral expression, incorporating the expertise of researchers into the classification process. However, a sizeable, manually-annotated data set is required to adequately learn and reproduce human rater annotations [8, 16–19]. A potentially more serious issue, reproducibility - between and within research groups - is known to suffer as the annotation process is prone to inherent biases and rater fatigue [6, 20]. Furthermore, as investigators aim to untangle a more complete and detailed behavioral repertoire, supervised algorithms are unable to generate new insights that build upon what they have already found or to match their subjective definitions of a given behavior across researchers. Related to this, these models often fail to provide a concise account of the quantitative basis for the learned decision boundaries - instead relying upon qualitative descriptions from the annotator’s intuitive reference frame [21]. Consequently, reproduction and comparison of the classification process can become a matter of subjective re-interpretation and escalating inter-rater variability.

The alternative approach is agnostic to experimenter definitions, focusing instead on uncovering the conserved spatiotemporal structures within pose dynamics [6]. These unsupervised models can find known and hidden behavioral expressions without the influence of human annotation. Unsupervised pattern discovery can be directly applied to provide behavioral expressions with temporal resolution beyond human ability into sub-second components and distinct sub-actions with high sensitivity [9–11, 22–24]. This major benefit is also its key drawback: the algorithm can only identify patterns that are statistically obvious given the input features provided - i.e., rare events or more subjective distinctions between behaviors will not be identified as unique clusters. Recent approaches tackle this issue by using additional metadata of animals in order to balance subsample the rare events or subtypes. [22]. However, this evidently requires either predefined categories or trial-and-error to optimize desired outcome performance. Thus, behavioral expressions that are evident to the experimenter - but are not readily discerned statistically - often cannot be reconstructed. Additionally, identified behavioral patterns are often assigned semantic names that may obscure more complex underlying feature statistics if used without proper validation. Conversely, behavioral patterns may be discovered that do not conveniently fit traditional nomenclature, particularly at temporal scales beyond the typical spectrum. To accommodate these cases, researchers often utilize token names (e.g., Grooming Sub-type A or Motif 1) that are difficult to interpret.

Summing up, supervised and unsupervised methods are prone to focus on different feature-integration approaches to segment behavior. As a result, each fails to combine informed analysis and efficient pattern discovery within the same workflow. An ideal solution would be able to reproduce an expert researcher’s informed annotations and translate them into a transparent, reproducible format. In addition, the researcher would be able to engage the power of agnostic discovery within the same framework - uncovering conserved movement patterns to facilitate deeper behavioral understanding.

Here, we present A-SOiD, an active supervised learning platform that incorporates unsupervised discovery of spatiotemporal movement patterns. A-SOiD outperforms traditional supervised methods in accuracy and greatly reduced training costs. By automatically balancing annotation sets, A-SOiD reduced the amount of annotations required by *≈* 90%. Because A-SOiD requires only a very small amount of input data to reproduce annotations, we were able to expand an initial set of behavior categories to include newly discovered behaviors autonomously with high predictive performance. It also provides an optional entry point for discovery to expand the classification annotation set through unsupervised segmentation of a selected behavior. To test its performance, we applied A-SOiD to two benchmark data sets. First, we applied it to social interactions in rodents that are complex and nuanced. While supervised approaches can be trained to identify select interactions, there are still substantial limitations on the specificity of behavioral segmentation - even with large training data sets. We investigated the sizeable human-annotated data set of social behavior in mice (CalMS21) [25] with A-SOiD. The dataset comprises not only a set of pose estimations and human annotations, but is also extensively described by the authors, including factors such as inter-rater variability, data distribution, their own solution [6]. We next extended the range of detected canonical social behaviors. This process should also not occlude the functional rationale behind newly found components. Notably, in the field of machine learning there are already developments striving to replace black-box approaches for better *post hoc* explainability [21, 26–28]. Moreover, we benchmarked against two popular supervised behavioral classification algorithms as well as public entries. Next, we addressed the current lack of supervised behavioral segmentation methods for both 3D pose estimation and for non-human primates, despite rapid growth in these fields [7, 29], especially concerning unsupervised approaches [24, 30, 31]. To demonstrate the application flexibility of A-SOiD, we analyzed a previously unpublished non-human primate three-dimensional pose data set. Finally, to facilitate A-SOiD’s use in the rapid segmentation of a behavioral repertoire, we packaged the platform into an intuitive open-source graphical user interface that can be used without prior coding experience (Supp. Fig. 1).

## Results

### User definitions of complex behavior are not readily identified using unsupervised approaches

Experimenters often focus their initial behavioral quantification on a few selected behaviors of interest, with the potential to expand their analysis by exploring the inherent data structure (unsupervised). This approach assumes that human definitions will be self-evident from the given data representation and that further explorations can be directly aligned with previous findings. Therefore, we first investigated whether an unsupervised classification approach that could be used to explore the data would also be able to reproduce human annotations given a large benchmark data set of socially-interacting rodents.

The CalMS21 data set is a large social behavior benchmark data set consisting of annotations from four expert human raters for three distinct behavioral categories (“attack”, “investigation” and “mount”; see Methods) were annotated (Fig. 1a-b, see Methods and [6, 25]). The data set also provides pose estimation data of the two socially-interacting mice (Fig. 1a). To extract a single, homogenized set of annotations, we divided the behavioral classifications into non-overlapping segments. For this, we examined the distribution of bout lengths across annotations and found that a period of 400 ms (12 frames 10%tile, Fig. 1c, Supp. Fig. 1) would be sufficient to resolve behavioral changes across annotations. Given that 12 frames could contain more than 1 type of annotation, we examined the annotation pattern in depth. We found that *≈* 92% of the 400 ms non-overlapped segments contained only a single behavioral annotation. The remaining *≈* 8% had at least 2 types of annotation within the same segments. Further examination of the remaining *≈* 8% cases revealed that a predominant annotation dominated the segments, thus, a tie-breaker was rarely required. Our results suggest that, compared to frame-wise annotation, there is minimal loss of information upon downsampling of annotations to 400 ms.

**Figure 1:**
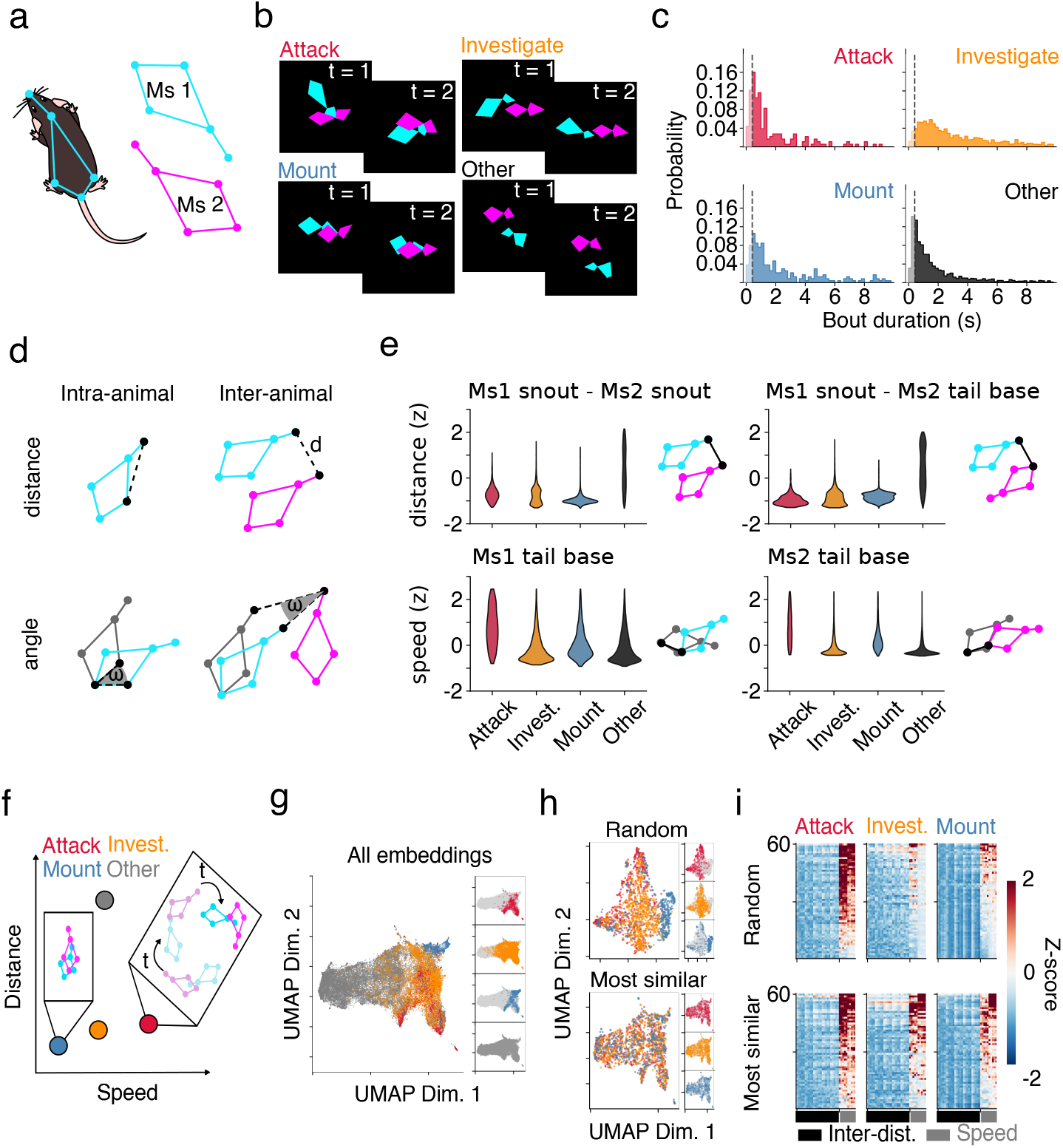
Human annotations of social interactions cannot be easily represented in unsupervised embeddings. a) Schematic representation of body positions on two mice (Ms1: cyan; Ms2: magenta) taken from the CalMS21 data set. b) Annotated example frames from the CalMS21 training data set (Red: “attack”; Orange: “investigation”; Blue: “mount”; Black: “other”). Behavioral color-code to be maintained throughout manuscript. c) Histograms showing the distribution of annotated frames before a transition into a different behavior occurs. Dashed lines indicate 400 ms (12 frames) that was used to integrate our features over, temporally. Session n=70. d) Intra-animal and inter-animal distances (top), and respective angular changes (bottom) are calculated for all combinations. e) Example feature distributions across annotated behaviors (top: snout-to-snout and tail base-to-tail base inter-animal distance, bottom: sample-to-sample tail base speed of Mouse1 and Mouse2). f) Schematic representation of unique composite feature distribution for each behavioral expression. g) UMAP embeddings of composite features, colored by annotations. h) UMAP embeddings using a random subset (top) and using the same count but a more similar subset. i) 2D-histogram showing normalized feature values (z-scored) across features (columns) and selected samples (rows) of all three classes (“attack”, “investigation”, “mount”). Here we show a subset (n=60) representing (h) for ease of visualization.

To capture the spatiotemporal pose relationships of the annotated social interactions, we extracted features (see Methods) from both the individual animals (intra-animal; Fig. 1d, left) and the multi-animal interactions (inter-animal; Fig. 1d, right). Notably, we observed that the distribution of single features was already indicative of the human annotations, e.g., mounting is characterized by a small inter-animal snout-snout distance while attacking typically possesses a greater speed (Fig. 1e, top-left). However, significant overlap can occur (compare Fig. 1e, bottom-left), which necessitates the evaluation of the composite feature distributions to fully represent a behavioral class (Fig. 1f).

With this realization, we next explored whether a popular unsupervised approach [9] (see Methods, Supp. Fig. 2d-f) could resolve the human annotation space given the extracted features (Fig. 1g). In the feature distribution (Fig. 1e), individual behaviors could be largely separated by a composite of several features. If the human annotations can be readily described by the selected features (Supp. Fig. 2), a high-dimensional representation of all points in the feature space or its low-dimensional embedding (UMAP) should provide distinct clusters that readily map onto each group (e.g., all examples of “attack” in the same area (high speed, low distance), whereas “mount” is in a different region (low speed, low distance); Fig. 1f, Supp. Fig. 2a-c). While an unsupervised approach has been proven successful for extracting common, single-animal behaviors [9–11, 22, 24, 32], when applied to the social interactions of the CalMS21 data set, we found that human annotation cannot be reliably represented as clusters of spatiotemporal pose relationships (Fig. 1g, Supp. Fig. 2d-f). Given the characteristics of unsupervised approaches, a potential problem is the highly unbalanced distribution of data in its raw state. The majority of annotations are designated as “other” (n=26409, see Methods), while all remaining behaviors are only represented by a substantially smaller amount of examples (“attack”: n=1188; “investigation”: n=12300, “mount”: n=2378). The clustering algorithm is inclined to emphasize the differences between data points in “other”, and consequently will overlook the differences in smaller classes. As such, we see that the “other” group spans the entire embedded space (Fig. 1g, grey), encompassing all other classes within. It is important to note that this bottom-up approach is agnostic to human classification and considers all points as equally important. The top-down human annotations were only added to the embeddings in order to visualize the lack of separation (Fig. 1g).

We further investigated whether a data set [25] comprised of a more balanced sample of all classes that contained clear definitions (“attack”, “investigation” and “mount”, see Methods) would yield better results in unsupervised embedding. Notably, we require human annotations to curate a more balanced sample of all classes. This would not be feasible on a completely unlabeled data set. First, we selected a random sample (n=580, 872, 581 for “attack”, “investigation”, and “mount”, respectively) and embedded them together to determine how an unsupervised approach would perform. While this embedding yielded better reproduction of the desired annotations, there appear to be subgroups within each annotation. Consequently, this approach does not resolve the entirety of the human annotation spectrum (Fig. 1h,i, top).

Next, while maintaining the same number of samples from the three classes, we selected those that appear most similar amongst the three annotation types. We discovered that there are a select group of samples that rarely differ in features (Fig. 1h,i, bottom). One possible explanation is that these samples represent transitions between two behaviors and could be labeled as either or - i.e., where does a rater decide when a behavior stops and a new one starts? We first examined the preceding behavior for each behavior that the algorithm deemed uncertain and found that when examining the subset of samples that are most similar in feature space (the “hardest to predict”), the consistency of annotations between the prior frame to the current frame (transition) is significantly lower than a random subset of samples (Supp. Fig. 3b). One key finding is that the rater’s decision at which frame the animal goes from “investigation” to “mount” is inconsistent, as there is a higher rate of “investigation” to “mount” transitions and vice versa (Supp. Fig. 3a). As expected, this is in line with the type of predictions errors by baseline models from Sun et al. [25].

Moreover, when investigating whether our classifications show a bias towards transitions “out of” a behavior or transitions “into” one, we found that when considering the samples that are the most similar, overall, the consistency of annotations between the current frame and the next frame is similar to transitioning “into” a behavior (Supp. Fig. 3d). More specifically, we found that, for the annotations in the used CalMS21 data, the particular inconsistency varies between behaviors for “into” vs “out of” transitions of the “investigation” class. For example, while the rater was less consistent with labeling a transition from “mount” into “investigation”, the inconsistency was higher for transitions out of “investigation” into other (Supp. Fig. 3a, c). Again, in line with prediction error types made by baseline models from Sun et al. [25].

While a purely data-driven approach such as unsupervised classification is able to identify behavioral clusters in underlying data [9, 11, 22, 24], it would require extensive optimizations and therefore is not the “go-to” solution to closely reproduce human annotations in this social interaction data set (see also Supp. Fig. 2d-f).

However, this interrogation of the data structure revealed key inroads to improved segmentation. Anticipating the challenges of an unbalanced data set and problematic edge scenarios, we focused on developing a solution that would automatically balance the training data and integrate human annotations in a data-efficient manner. Finally, we embraced the possibility that a data-driven approach, such as unsupervised clustering, may perform better when applied to the restricted, segmented classes - and allow further discovery of conserved patterns.

### Active learning automatically balances training data and integrates human expertise with high performance

We developed A-SOiD - an intuitive GUI pipeline (Fig. 2a; Supp. Fig. 1) that includes both active learning and directed unsupervised behavioral segmentation. A-SOiD makes use of an iterative, active learning paradigm that selectively trains on low-confidence examples to improve classification robustness. Rather than unselectively feeding in additional human annotations, A-SOiD queues a subset of low-confidence predictions from the rest of the training using each previous classifier iteration. By focusing on the potentially problematic edge cases (Fig. 1h, i, bottom, Supp. Fig. 3), the algorithm greatly reduces the overall number of annotated frames required (Fig. 2a, steps 1-3). Beyond initial annotations, A-SOiD provides users with an unsupervised clustering algorithm [9] to explore and further subdivide existing annotations (Fig. 2a, step 4).

**Figure 2:**
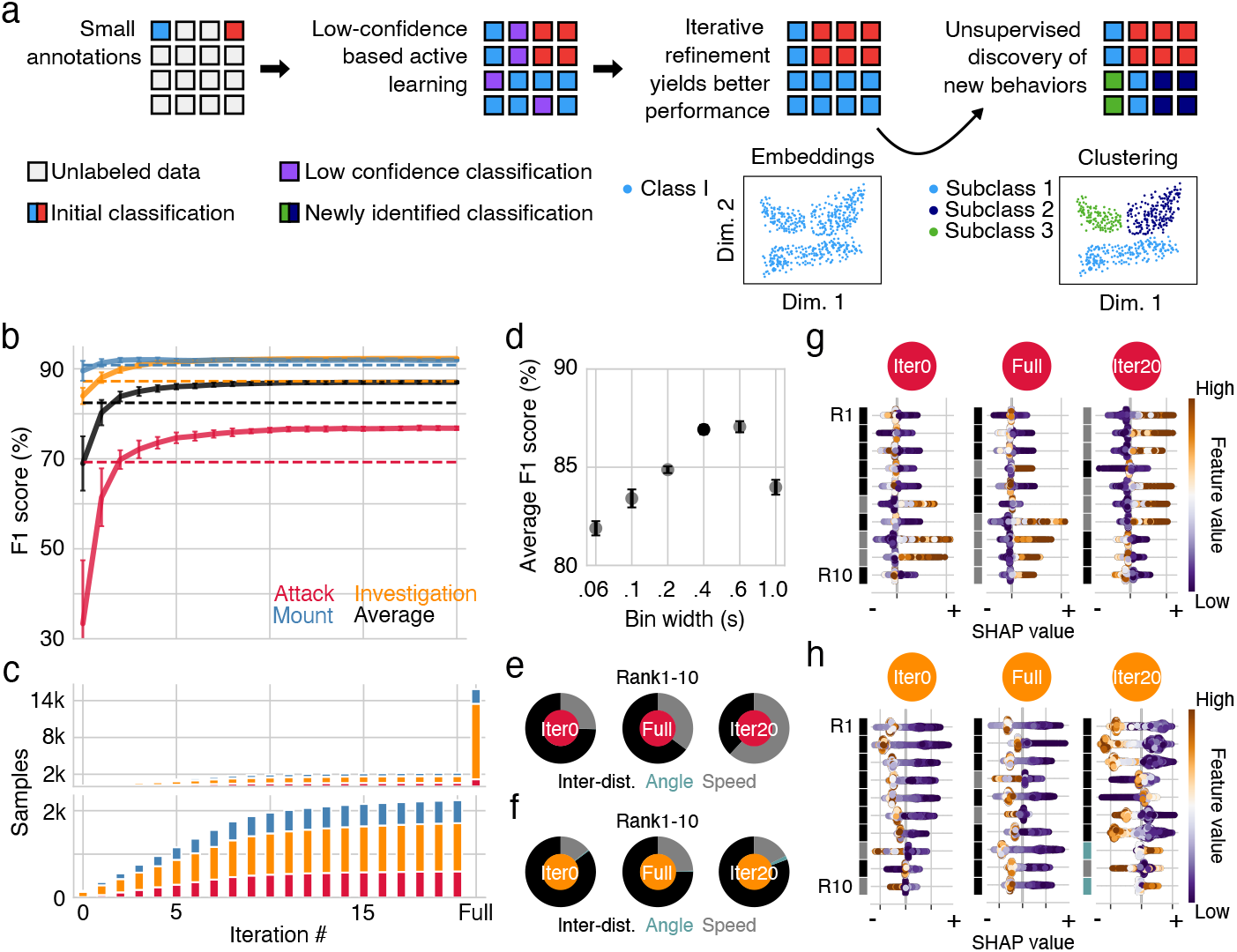
Active learning improves data efficiency and overall performance via different feature weighting. a) The A-SOiD pipeline initializes a classifier with minimal training annotations (step 1). Next, low-confidence predictions on the remaining training annotations are intelligently queued up for the next training iteration (step 2). Step 2 reiterates until there are no remaining low-confidence training samples (step 3). Lastly, A-SOiD can expand the annotation set by deploying directed, unsupervised pattern discovery (step 4). Class I is subdivided into 3 subclasses through B-SOiD. b) A-SOiD performance (F1 score, 20 cross-validations, top) on held-out test data is plotted against active learning iterations. Dashed lines demonstrate performance using full training annotations at once. Red: “at- tack”; Orange: “investigation”; Blue: “mount”; Black: Unweighted class average. Error bars represent *±* standard deviation across 20 seeds (see Methods). c) Stacked bar graph of the number of training annotations is plotted against active learning iterations. The right-most bar represents the full annotation count. Red: “attack”; Orange: “investigation”; Blue: “mount”. Bars represent average training samples across 20 seeds. d) Unweighted average performance on held-out test data is plotted against 6 different window sizes (sub-second to a second) for feature binning. e) Pie charts of the most impactful feature components (distance, angle, or speed) where 20 iterations of active learning (right) corrected “attack” misclassification of either using complete annotations at once (middle) or prior to undergoing active learning (left). Black: inter-distance; Teal: Angular change; Gray: Speed. f) Pie chart of the most impactful feature components (distance, angle, or speed) where 20 iterations of active learning (right) corrected “investigation” misclassification of either using complete annotations at once (middle) or prior to undergoing active learning (left). Black: inter-distance; Teal: Angular change; Gray: Speed. g) For the top 10 most impactful features (rows, Rank 1 (R1) to Rank 10 (R10)) where “attack” was correctly classified in iteration 20 (right), but not otherwise (left/middle), the color-coded feature value is plotted against the SHAP value, for “attack” specifically. Note that positive SHAP values (to the right) will succeed in reproducing the classifier’s prediction for “attack”. h) For the top 10 most impactful features (rows, Rank 1 (R1) to Rank 10 (R10)) where “investigation” was correctly classified in iteration 20 (right), but not otherwise (left/middle), the color-coded feature value is plotted against the SHAP value, for “investigation” specifically. Note that negative SHAP values (to the left) fail to reproduce the classifier’s prediction for “attack”.

To directly overcome the bottleneck of supervised approaches requiring massive training data sets while reducing the possibility of de-emphasizing the smaller annotations like “attack”, we first initialized A-SOiD by providing a mere 1% of samples that were randomly selected from each of the three annotations. We maintained the relative proportions when initializing to maintain a relatively similar feature distribution to the entire pool. Not surprisingly, the resulting predictions underperform compared to using the entire data set (Fig. 2b, iteration 0). Next, a selected subsample of low confidence predictions on the remaining training data (i.e., most similar across classes, Fig. 1h,i bottom) by the initial classifier (iteration 0) is refined by extracting annotations from the remaining training labels or alternatively by prompting human refinement. In subsequent iterations, the latest classifier is used to identify new low-confidence predictions and to refine them for the training of the following iteration. This establishes an iterative active learning scheme that focuses on clarifying annotation preference only for examples that lie at the decision boundary between classes (Supp. Fig. 3). With this approach to refinement, performance on a completely unseen, held out test data set improves beyond using the entire available training data with just 10% of the labels necessary, or fewer than 1000 samples per class (Fig. 2b, the dashed line represents performance using all training data, black indicates average across all three classes). A-SOiD performance reached 0.870 *±* 0.002 macro average f1 score and 0.918 *±* 0.001 MAP with 20 cross-validation runs - on par with *Top-1* performance in the MABe 2021 Task1 Challenge [25], and without the use of additional unlabeled data set. Note that, at no point in time was the test set used to improve algorithm performance. The performance plateau is paralleled by a drop in additional samples per iteration as the quantity of low-confidence samples sharply drops. Additionally, we observed that the number of training samples per behavioral expression became more balanced out over active learning iterations, even when the total available counts, and subsequently our initialized classifier inputs, were largely biased towards one behavior expression (“investigation”; Fig 2c, orange of full dataset). Therefore, these benchmark data suggest that even if A-SOiD starts from scratch with a small, random sample of frames, performance quickly exceeds that of alternative classifiers trained on the all training data at once by nearly an order of magnitude less training samples. Furthermore, we examined the feature window size’s effect on performance and found that the 10 percentile (or 400ms) window size best captures these behavior dynamics (Fig. 2d). Upon reaching 20 iterations of active learning, A-SOiD outperformed predictions using the full training data (Fig. 2b solid line versus dashed line). To understand why a correct or incorrect prediction might be made, we used an explainability metric to assess the impact of individual features in the decision process using SHAP analysis (SHapley Additive exPlanations) [26, 27]. SHAP is a method that infers the credit assigned to the individual feature underlying the performance of an algorithm using a game theoretic approach (see Goodwin [21] and Mathis [28] for discussion of the value and comparison of trade-offs for different explaniability metrics). Looking at the category with the most mistakes using the full model, “attack”, SHAP identified a larger emphasis on speed-related components (**L**, see methods) in the ten most impactful features (gray component in right and middle plots Fig. 2e). This bias towards speed features was not observed prior to learning iterations (Iter0, Fig. 2e, left).

When examining a particular instance where the 20-iteration model correctly chose “attack” and the full model did not, we found that the top-ranked speed-related components drove the “attack” probability much higher while decreasing the “investigation” probability, which did not occur when either using the full data set or prior to active learning (Supp. Fig. 4a, c, e). This increase in the importance assigned to speed is somewhat intuitive; it is the increase of speed of movement that largely differentiates attacking from other social interactions (Fig. 2g, right). In the next most error-prone classification, “investigation”, our SHAP analysis pointed to the incorporation of body angle - specifically, (Θ, see methods) as a key difference between iteration 20 and either the full data set or pre-learning (2f, teal). When examining an instance when the 20-iteration model was correct but the full model was not, we found that the top ranked angular components drove the “investigation” probability higher while decreasing the “other” probability (Supp. Fig. 4b, d, f).

### Benchmarking performance across supervised segmentation methods

To quantify the performance of A-SOiD against the state-of-the-art in supervised behavioral segmentation, we compared the performance of A-SOID to other approaches. The CalMS21 data set was created to competitively evaluate different approaches during a machine learning challenge [6]. Notably, even when allotted 10 times less training data than was available, A-SOiD outperformed the best solution in the competition by a small margin[6] (Fig. 3b). Using this same benchmarking dataset, we compared A-SOiD’s performance to two popular supervised solutions, SimBA [8] and DeepEthogram (DEG) [33]. Various performance metrics indicated that A-SOiD matched human raters best (unweighted macro F1: Fig. 3b; weighted macro F1: Supp. Fig. 5c; micro F1: Supp. Fig. 5b.). Since the CalMS21 dataset described the “other” as not a behavioral category, we showed the prediction limited to “attack”, “investigation”, and “mount” (Fig. 3a, prediction including “other” in Supp. Fig. 5a). A-SOiD outperformed these methods across all three annotated behaviors (Fig. 3c, Supp. Fig. 5f). We observed that human raters and A-SOiD demonstrated a similar ratio of “attack”, “investigation”, and “mount” predictions in the data - while SimBA and DEG overrepresented its most common category, respectively (Fig. 3f, Supp Fig. 5d, e). DEG differs from SimBA in that it allows for a background class rather than assigning a class. The background class, in other words, is a behavior that could be anything other than the three classes. On the other hand, SimBA does not allow for such an option, requiring a fourth binary classifier that recognizes “other”. A-SOiD outperformed both SimBA and DEG because there were fewer false positives in either “other” or “investigation” (Fig. 3c bottom). Overall, these results demonstrated that A-SOiD performed better by reducing the bias towards dominant classes through feeding in a more balanced multi-class training set.

**Figure 3:**
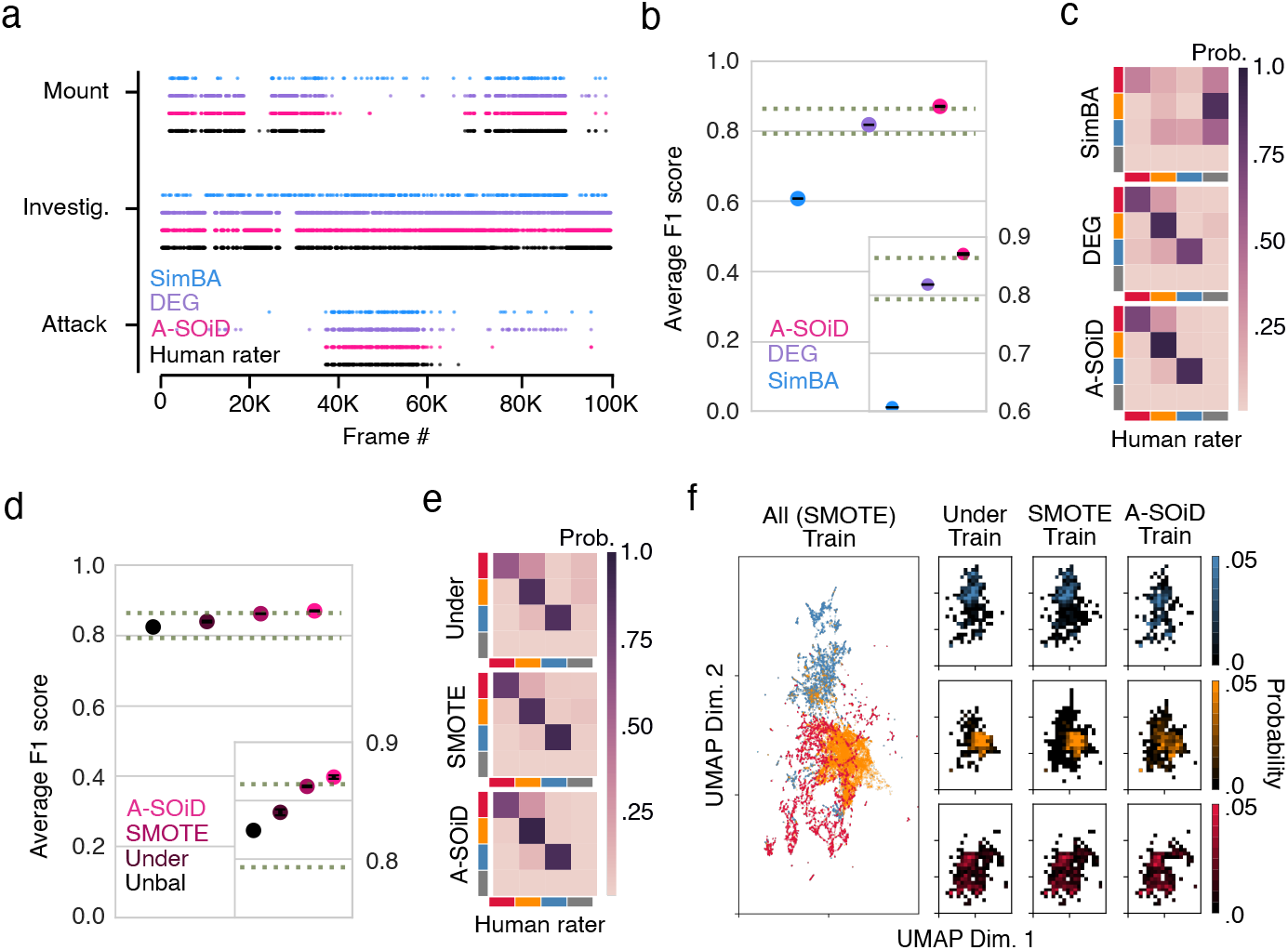
Benchmark results against state-of-the-art supervised classification methods. Predicted ethogram using SimBA (blue), DeepEthogram (DEG, purple), and A-SOiD (pink) against target human annotation (black). Every fiftieth frame is shown to allow for better visualization. Session n=19. b) Scatter plot of unweighted mean F1 scores across all behaviors using SimBA (blue, left), DEG (purple, middle), and A-SOiD (pink, right). The top dotted line represents the winner across 48 entries to challenge, while the bottom dotted line represents the MARS baseline model presented in [6]. Inlet scatter plot highlights the performance difference. Error bars represent *±* standard deviations across 20 seeds (see Methods). c) Confusion matrices where prediction mistakes were being made for SimBA (top), DEG (middle), and A-SOiD (bottom). Darker shades along the diagonal indicate better algorithm performance in matching the human rater target. d) Scatter plot of unweighted mean F1 scores across all behaviors using complete annotation but unbalanced (black, left most), random under-sampled but balanced (brown, second to the left), SMOTE oversampled and balanced (maroon, second to the right), and A-SOiD (pink, right most). The top dotted line represents the winner across 48 entries to challenge, while the bottom dotted line represents the MARS baseline model presented in [6]. Inlet scatter plot highlights the performance difference. Error bars represent *±* standard deviations across 20 seeds. e) Confusion matrices where prediction mistakes were being made for random under-sampled but balanced (top), SMOTE oversampled and balanced (middle), and A-SOiD (bottom). Darker shades along the diagonal indicate better algorithm performance in matching the human rater target. f) Low dimensional embedding of SMOTE oversampled training features (left). Within the embedding, 2D histograms per annotation (“mount”: top; “investigation”: middle; “attack”: bottom) are plotted for random under-sampled training data (left), SMOTE oversampled training data (middle), and A-SOiD (right) intelligently recruited training data.

**Figure 5:**
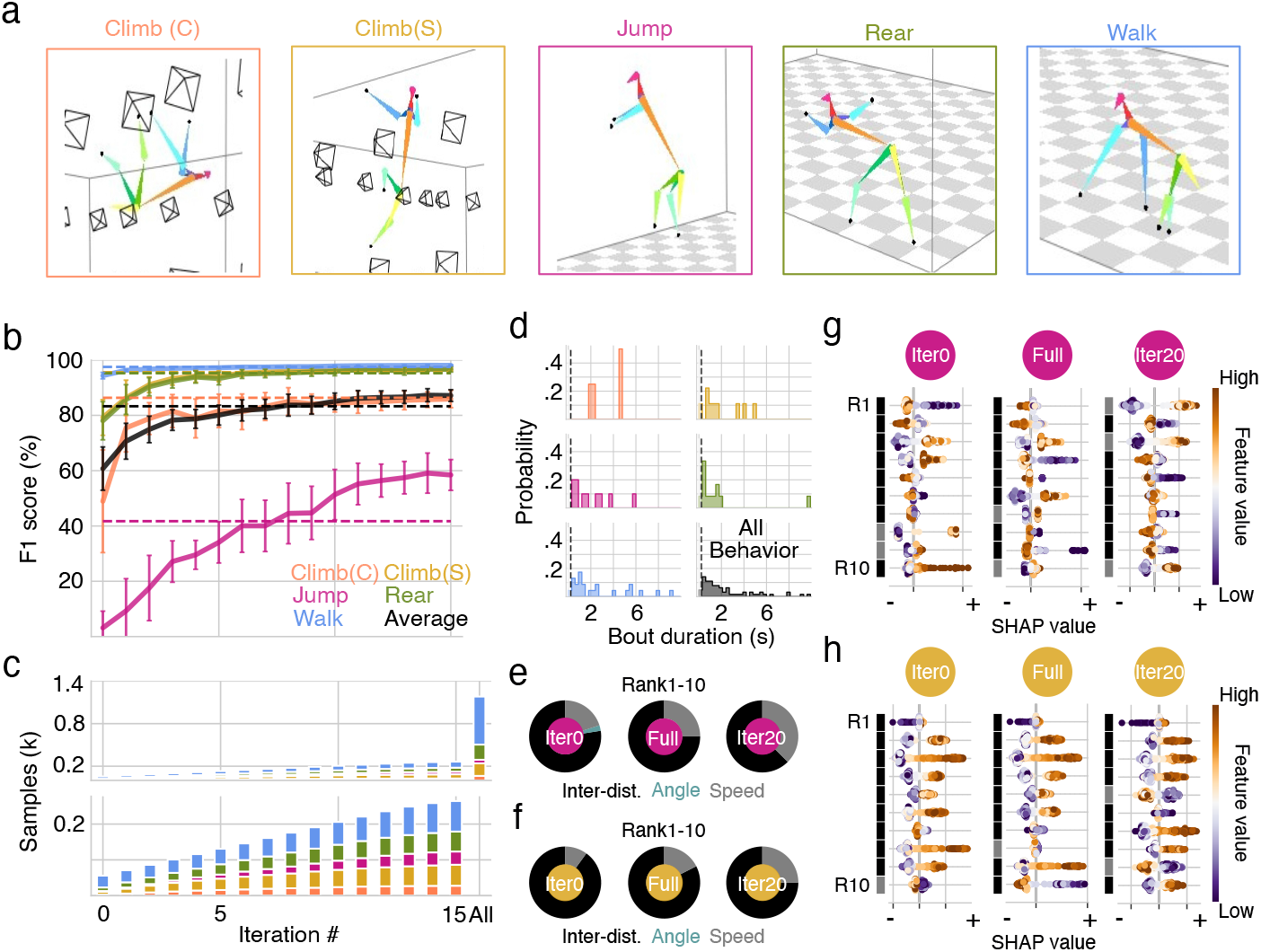
Efficient segmentation of monkey behavioral repertoire a) Representative frame examples for each annotated behavior. Images are reconstructed by OpenMonkeyStudio (3D pose). b) A-SOiD performance (F1 score, 20 cross-validations, top) on held-out test data is plotted against active learning iterations. Dashed lines demonstrate performance using full training annotations at once. Orange: “ceiling climb” (Climb C); Yellow: “sidewall climb” (Climb S); Pink: “‘jump”; Green: “rear”; Blue: “walk”; Black: Unweighted class average. Error bars represent *±* standard deviation across 20 seeds (see Methods). c) Stacked bar graph of the number of training annotations is plotted against active learning iterations. The right-most bar represents the full annotation count. Orange: “ceiling climb” (Climb C); Yellow: “sidewall climb” (Climb S); Pink: “jump”; Green: “rear”; Blue: “walk”; Bars represent average training samples across 20 seeds. d) Histograms showing the distribution of annotated frames before a transition into a different behavior occurs. Dashed lines indicate 200 ms (6 frames) that was used to integrate our features over, temporally. Session n=1. e) Pie charts of the most impactful feature components (distance, angle, or speed) where 20 iterations of active learning (right) corrected “jump” misclassification of either using complete annotations at once (middle) or prior to undergoing active learning (left). Black: inter-distance; Teal: Angular change; Gray: Speed. f) Pie chart of the most impactful feature components (distance, angle, or speed) where 20 iterations of active learning (right) corrected “climb (S)” misclassification of either using complete annotations at once (middle) or prior to undergoing active learning (left). Black: inter-distance; Teal: Angular change; Gray: Speed. g) For the top 10 most impactful features (rows, Rank 1 (R1) to Rank 10 (R10)) where “jump” was correctly classified in iteration 20 (right), but not other-wise (left/middle), the color-coded feature value is plotted against SHAP value. Note that positive the SHAP values (to the right) will succeed in reproducing the classifier’s prediction for that class. h) For the top 10 most impactful features (rows, Rank 1 (R1) to Rank 10 (R10)) where ‘climb (S)” was correctly classified in iteration 20 (right), but not other-wise (left/middle), the color-coded feature value is plotted against the SHAP value. Note that negative SHAP values (to the left) fail to reproduce the classifier’s prediction for that class.

We next investigated whether there are additional benefits beyond multi-class annotation balance when using A-SOiD. Synthetic minority over-sampling technique (SMOTE) [34] has been proven effective in data balance using underlying feature distribution. To compare and contrast A-SOiD’s performance against existing data balance methods, we benchmarked against unbalanced (unbal), random under-sampling (under), and SMOTE. We found that data balance improves performance regardless of method (Fig. 3d). However, A-SOiD still outperforms state-of-the-art data balancing technique (again, with much less training samples; Fig. 3d). Interestingly, the largest class: “investigation” was the main difference between A-SOiD and these two data balancing techniques (Fig. 3e, Supp. Fig. 5g). This is consistent with the described pitfalls of using SMOTE with high-dimensional datasets, where, ultimately, the majority class (the only class that does not get synthetic data points) gets overshadowed, leading to classification biases towards minority classes at the expense of the majority class [35, 36]. Given this line of reasoning, we explored the class-specific means in the CALMS21 dataset. To examine class-specific means, we first embedded the SMOTE up-sampled features. In this fixed manifold, we can examine the samples that were used in random under-sampling (under), SMOTE, and A-SOiD all in the same axes (Fig. 3f). What became evident was the reduced density in A-SOiD’s “investigation” (Fig. 3f, left most orange), whereas SMOTE does not alter the class-specific means from random undersampling (Fig. 3f), in line with the previous literature describing SMOTE’s limitation [35].

### Expanding segmentation utility through fast and efficient sub-pattern discovery

While A-SOiD’s active-learning component allows the reproduction of human annotations in a highly efficient manner, it lacks the ability to discover conserved patterns of behavior and thereby limits researchers to their initial set of behaviors. This limited set may be due to a lack of comprehension of the behavioral repertoire during annotation or difficulties in robustly defining rare or nuanced sub-actions. For example, the CalMS21 data set does not inherently differentiate between the several known “investigation” types. As such, investigative behavior sub-types are hidden and dispersed within the “investigation” class (Fig. 4). However, it is often desirable to be able to split high-level behaviors into conserved sub-actions. Specifically, in social interactions encompassing both male and female mice, “anogenital investigation” is a primary behavioral expression when mice engage in olfactory investigation. Quantification of such behavior can therefore serve as a key identifier for social recognition, including habituation and discrimination [37, 38] (https://mousebehavior.org/investigate-anogenital/) which in turn indicates a branching point in behavioral strategies depending on the specific outcome.

**Figure 4:**
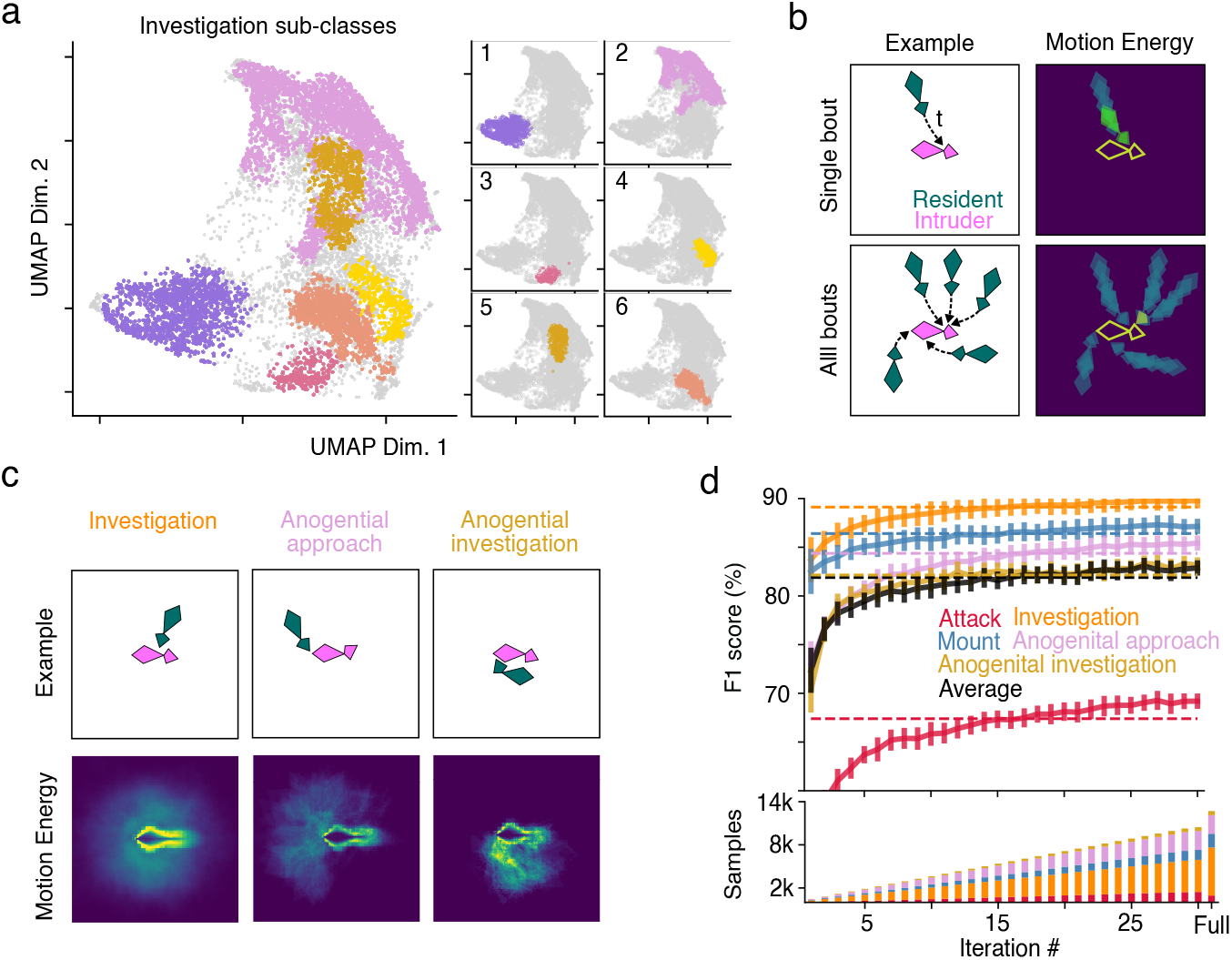
Unsupervised Clustering can be used to discover and integrate novel behavioral expressions in previously unspecific data. a) UMAP embeddings followed by HDBSCAN clustering (see Methods) of only the “investigation” class revealed 6 behavioral sub-classes. b) Schematic representation of motion energy in a single bout (top) and average across all bouts (bottom). c) Representative examples (top row) and average motion energy (bottom row) of the “investigation” class, including two sub-types that closely align with the heuristic identification of “anogenital investigation” (Suppl. Fig. 5). d) With new sub-classes redefined within the existing training data set, A-SOiD (F1 score, 20 cross-validations, top) outperforms a classifier trained on the full data at once (dashed lines). Note that in this case, we are only considering a subset of the CalMS21 training data (80%/20% split within the CalMS21 training data described throughout the manuscript) to allow testing on the remaining data. This performance increase is achieved through less data (bottom). Red: ”attack”; Orange: ”investigation”; Blue: ”mount”; Pink: “anogenital approach” (sub-class 2); Gold: “anogenital investigation” (sub-class 5); Black: average across classes. error bars represent *±* standard deviation across 20 seeds (see Methods). Bars represent average training samples across 20 seeds.

Therefore, for the directed discovery of hidden sub-behaviors, we included an optional unsupervised classification step within previously annotated behaviors (top-down). This allowed us to preselect behaviors of interest for further subdivision and discover conserved investigative sub-types when applied to the investigation class. Unsupervised embedding and clustering of the “investigation” class separately from the full data set revealed several sub-classes (Fig. 4a, see Methods). Of these, two clusters appear to be “investigation” specifically at the anogenital areas. More specifically, one of the sub-classes consists of the resident mouse approaching and directing its “investigation” at the anogenital area of the intruder mouse from behind (sub-class 2, “anogenital approach”, n=3234) - and the other of the resident mouse investigating the anogenital area of the other mouse while already in close proximity (sub-class 5, “anogenital investigation”, n=706; http://mousebehavior.org/ethogram-index/; Fig. 4c). To allow a direct comparison between unsupervised clusters and the heuristic criterion that the snout of one or both mice must be close to the tail base of the other, we annotated all data points within the “investigation” class in which the distance between the resident’s snout and the intruder’s tail base was lower than a manually set threshold (15 pixels; Supp. Fig. 6). A comparison revealed an extensive overlap between the unsupervised clusters of sub-class 2 and sub-class 5 with the top-down, manually selected feature space.

We then computed the motion energy (see Methods) for each cluster. Motion energy analysis results in single images that average the motion spectrum relative to an individual, aligned animal (Fig. 4b). Consequently, conserved behaviors with repeated distinct movements result in a bright, clearly defined image, while behaviors that include divergent movement patterns appear darker and blurry (Fig. 4b). In both example clusters, the average resident snout’s motion energy is concentrated close to the anogenital region of its con-specific, unlike the widely distributed motion energy found in the “investigation” class (Fig. 4c). Specifically, the sub-class 2 appears restricted to motion behind the centered animal, i.e., a targeted approach to the anogenital area, “anogenital approach”, while sub-class 5 consists of paralleled resident/intruder anogenital investigations, “anogenital investigation” (Fig. 4b). We confirmed the motion energy inspection by extracting example episodes (Supp. Video 1) of the found behavioral subtypes.

To test A-SOiD’s ability to integrate discovered sub-types into our classifier, we provided additional labels for sub-class 2 (“anogenital approach”) and sub-class 5 (“anogenital investigation”) by splitting them from “investigation”, while all remaining clusters remain in the original class “investigation” (Fig. 4a). We then retrained a classifier using the expanded annotation set. Since we only clustered the provided training set from the CalMS21 data set, we decided to split the training set into a test and train set, which resulted in a reduced overall performance in this particular section compared to active learning using the full training set (compare with Fig. 2b). Similar to the active learning performance described in (Fig. 2b), we found that after 30 iterations, we reached higher predictive performance across all categories, while “attack” still improved the most (Fig. 4d). Taken together, these results demonstrate the possibility of iterating between supervised and unsupervised classification.

### A-SOiD demonstrates improved performance regardless of species or spatial dimensions

To demonstrate the flexibility of our approach, we applied A-SOiD to position information and human annotations of singly-housed non-human primates. Notably, pose information was computed using OpenMonkeyStudio [7], which provides 3D pose estimations. The video was manually annotated, separating the animal’s behavior into 5 categories: “walk”, “rear”, “jump”, “ceiling climb” (Climb C), and “side-wall climb” (Climb S), (Fig. 5a, and see Methods for a detailed description of our annotations). Next, we trained A-SOiD to reproduce the human rater annotations (n=64, 177, 50, 214, 676 for “ceiling climb”, “sidewall climb”, “jump”, “rear” and “walk” respectively). In this case, 15 active learning iterations were sufficient to reach plateau performance (Fig. 5b). Similar to the performance in the social mouse benchmarking, we found that A-SOiD automatically balanced the training set to include examples of classes that were initially underrepresented across iterations (“jump”, Fig. 5c). The overall predictive performance improved beyond the performance of a classifier trained on the full data all at once (compare Fig. 5b, dashed lines). Similar to the larger CALMS21 dataset, we examined the distribution of bout lengths across annotations and found that a period of 200 ms (6 frames 10%tile, Fig. 5d, left of dashed lines) would be sufficient to resolve behavioral changes across annotations. Upon reaching 15 iterations of active learning, A-SOiD outperformed predictions using the full training data (Fig. 5b, solid line versus dashed line). Again, to investigate why a correct or incorrect prediction might be made, we assessed the impact of individual features in the decision process using SHAP analysis. Looking at the category with the most mistakes using the full model, “jump”, SHAP identified a larger emphasis on speed-related components (**L**, see methods) in the ten most impactful features (gray component in right and middle plots Fig. 5e). This bias towards speed features was not observed prior to learning iterations (Iter0, Fig. 5e, left).

When examining a particular instance where the 20-iteration model correctly chose “jump” and the full model did not, we found that the top-ranked speed-related components drove the “jump” probability much higher while decreasing the “walk” probability, which did not occur when either using the full data set or prior to active learning (Supp. Fig. 6a, c, e). This increase in the importance assigned to speed is also intuitive; it is the increase in speed of movement that largely differentiates jumping from walking (Fig. 5g, right). This was also exemplified in the classification of “climb (S)” (5f, Supp. Fig. 6b, d, f).

Finally, we also assessed A-SOiD performance in a substantially larger human 3D-pose dataset [39] comprised of 11 social behaviors and 49 single-subject behaviors. There was a diverse array of complex human behaviors including varied object use (e.g. “taking off glasses”, “making a phone call”, “playing with phone”), different degrees of interpersonal contact (“patting”, “pushing”, or “punching”), and the expression of internal states (“cheer up”, “nausea”). Compared to the average performance using one-shot learning on the entire balanced dataset, A-SOiD yielded equivalent performance with 23% less training data (social; Supp. Fig. 10). This performance was also substantially better than that observed when using a subset of data with the equivalent number of training frames. Taken together, our results demonstrate A-SOiD’s flexibility across behavioral target types and numbers of targets.

## Discussion

As computational ethology explores the power of Big Data, the tools used to extract meaningful information must enable researchers to report transparent findings and allow generalizability across experiments. This is especially true in neuroethology, where variations in behavioral detection can fundamentally alter neural correlations or preclude their detection entirely [6, 9, 10, 20, 40]. To this extent, public benchmark data sets such as the social rodent data set CalMS21 [25] serve as a meaningful resource for developing and benchmarking new tools. We used this data set to develop a framework for the data-efficient integration of human-expert annotations and directed pattern discovery called A-SOiD. We then demonstrated the power of the approach, applying it to a completely unrelated data set - primate 3D pose estimation data.

Thus, A-SOiD facilitates building user-defined behavior analysis pipelines with high data efficiency. A-SOiD’s active learning framework resulted in a 10-fold decrease in required samples (total available labels=15866, used labels at performance peak=1866; see Supp. Table 2) to reproduce human annotations of socially active rodents with high performance (Fig. 2b). Even in the small non-human primate data set (n=1181 total labels), A-SOid robustly increased the performance of the low-frequency behavior (“jump”, n=50) by selectively training on low-confidence examples (Fig. 5b, see Supp. Table 3). Moreover, the resulting analysis of the model’s output can reach different levels of complexity depending on the research question. The unsupervised clustering of the embedded space of an annotation-based class was able to discover well-known behavioral phenotypes that were not the focus of the initial annotations and likely would not have been possible with a purely heuristic approach (see Fig. 4, Supp. Fig. 6).

**Table 2:** Actions included in the human data set (social)

| ID | description | ID | description |
| --- | --- | --- | --- |
| S01 | punching/slapping other person | S07 | giving something to other person |
| S02 | kicking other person | S08 | touch other person’s pocket |
| S03 | pushing other person | S09 | handshaking |
| S04 | pat on back of other person | S10 | walking towards each other |
| S05 | point finger at the other person | S11 | walking apart from each other |
| S06 | hugging other person |  |  |

### Quantifying and comparing classifier performance

An important unresolved issue in behavioral classification is the quantification of classification performance.

Often, the go-to approach is to directly evaluate the performance of a given approach by comparison to similar solutions in the field using performance metrics. However, while solutions often aim to solve the same general problem, a fair comparison is often difficult because each is showcased with in-house example data sets. Here, it is highly beneficial to work with open benchmark data sets such as CalMS21 to facilitate a fair comparison. In fact, we specifically chose to develop our solution with the data set because researchers would be able to directly compare the performance with other published solutions [6]. However, as this benchmark data set is relatively newly published, some state-of-the-art solutions have not been directly evaluated. To this extent, we expanded the available collection of comparable solutions with two additional state-of-the-art methods within the field (DeepEthogram and SimBA) [8, 33] by benchmarking them with CalMS21 data.

In our hands, A-SOiD reached higher performance than all other tested solutions (Fig. 3b) with 10 times less training data. Interestingly, a visual comparison of example frames reveals that the overall detection of behavior is similar across all methods (Fig. 3a), but A-SOiD manages to reduce false positive classifications of the dominant “investigation” behavior, of which helped the low-frequency “attack” and “mount” behaviors be more aligned to target human annotations (Fig. 3c, d). Our approach emphasized the importance of training data balance across behavioral classes and the possible gain from concentrating on the algorithm’s insecurities during training (Fig. 3b bottom). Moreover, in bench-marking against various algorithms that are available to the public, we sounded an alarm in studying behavioral bout duration for either having a biased prediction for “investigation” such as the use of DEG in our case and using potentially noisy pose data for classification.

In general, while all solutions aim to solve a supervised learning task, each approach has its own advantages and disadvantages, so that a comparison purely based on classification performance, even with a benchmark data set, is limited in its explanatory power. For example, while SimBA and A-SOiD use pose estimation data to extract movement features, DeepEthogram relies on raw video data and trains a set of three different neural networks to perform the same task. Consequently, the overall resource and time requirements to establish such an approach vary drastically. For example, the training of the DEG models required for the evaluation took approximately 22 hours on a high-performance workstation (264 Gb RAM, 24 Gb GPU), while A-SOiD can be run on new data on a standard laptop within an hour from installation. When using a Google Collab notebook (Free version, 12.7 Gb RAM) to provide standardization and an estimate of A-SOiD’s capability to be run on external servers, all 42,275 samples (CalMS21) could be processed in 7.63 minutes (458.34 +/- 5.81 sec, n = 3), including the entire active learning regime. Extrapolating from subsets of the CalMS21 data set (see Suppl. Fig. S8), 100 thousand samples would require 10.43 minutes, while one million samples would only require 53.24 minutes. Apart from this, DeepEthogram and other methods cannot parse multi-camera or 3D data pose data, making A-SOiD the only supervised method to use for nearly all primate and a rapidly increasing percentage of rodent studies.

Furthermore, while comparison between performance scores given a labeled data set is possible and there is a general consensus concerning the relative features of several behaviors, there is no ground truth to translate human expertise into operational definitions and vice versa without inevitable inaccuracies during reporting. In other words, when splitting behavioral subtypes within “investigation”, many may disagree or become biased by experience with the boundary between “anogenital approach” vs. “anogenital investigation”. Thus, there is a need for comparative methods that can be used to quantify algorithmic performance in a transparent and reproducible way. In this manuscript, we utilize two approaches that focus on the internal consistency of behavior classes themselves rather than an external, top-down rule.

Motion energy can quickly asses the differences between movement patterns (e.g., behavioral sub-types). Motion energy is an intuitive and informative way to generate a visual summary of the action within a found cluster [41]. In our hands, we utilized motion energy images to quickly differentiate sub-types of “anogenital investigation” (Fig. 4). Note, that comparative analysis can be done by analyzing the energetic variability within and across groups, providing a valuable statistic for cluster quality [9].

Another approach is using algorithmic explainability metrics, such as SHAP-based reporting [26, 27] of the underlying feature importance, which can help to share and compare conserved patterns across studies (for a review, see [21]). In this study, we employed SHAP analysis to investigate the algorithm’s learning process across multiple iterations during active learning and could identify that specific sets of features accounted for the increased performance of our classifiers. Once the algorithm is trained, these same values can be used to compare the classification reference frame across various models.

### Addressing variations in human annotations

Inconsistencies between human raters are a general issue of supervised approaches. Regarding the CalMS21 dataset, extensive analysis has been done adressing this issue [6]. However, for A-SOiD, we begin by selecting a small number of samples for training. Small training sets have the tendency to fail at representing real-world scenarios if they are not specifically selected (e.g., under-sampling from a bigger pool). As A-SOiD reproduces the human reference frame, biases or mistakes introduced by the annotations will consequently impact the final classification. Therefore, researchers utilizing A-SOiD on their own data can primarily counteract these issues by applying basic principles of data annotation such as multi-rater annotations [42, 43]. Additionally, our active learning strategy based on low-confidence examples emphasizes the majority of the sample representations, omitting rare mistakes and occasional variations. Importantly, when annotating behaviors, experimenters are predisposed to mirror the biased distribution of behaviors in the data set, thus producing an unbalanced training data set. To improve the performance of supervised algorithms, experimenters add additional frames, but this rarely overcomes the initial imbalance. However, the root of the problem is that classifier objective functions strive to improve overall performance. Although there are good solutions for binary imbalanced classifications [44–47], multi-class imbalanced classification is not as well developed [48, 49]. The main issue lies in the varying relationships between classes, as there could be two out of three classes that are balanced (both large), while the third one is small, or vice versa [50]. Our solution is to implement the intelligent selection of samples (active learning) within the already given annotations to reduce the bias towards larger classes. By starting with an absolutely minimal number of annotations, we let A-SOiD determine which samples are to be annotated. This approach effectively sparsifies representation while focusing on the outliers. While similar approaches have been previously utilized to allow the fast and efficient training of models in the field of pose estimation [1, 4] and behavior classification [19], they were integrated as implicit features to allow users the refinement of training data. In contrast, A-SOiD implemented active learning as an explicit core feature, and samples are selected automatically to be refined.

### Intuitive A-SOiD GUI for improved research integration

To improve the efficacy of this approach, A-SOiD comes as an app that can be installed on local computers without additional coding (Supp. Fig. 1). The app guides researchers through active learning and facilitates training high-performing classifiers based on an initially labeled data set. Researchers are then enabled to further refine their classifiers on unlabeled data using an intuitive interface in which low-confidence examples are presented for replay and evaluation. While the app is capable of reproducing the results from the CalMS21 data set reported in this study, we expanded its data importation capabilities to include the two most common open-source pose estimation solutions (DeepLabCut [1] and SLEAP [4]). Further, once a classifier reaches satisfactory performance levels, the app can be used to predict behaviors on novel data directly. The classifier can also be exported and deployed in custom analysis pipelines, including closed-loop experiments [12].

A-SOiD targets an unmet need in behavior analysis tools and provides researchers with an accessible, conjoined solution of supervised and unsupervised classification.

## Acknowledgements

We thank the labs of Drs. Jan Zimmermann and Benjamin Hayden at the University of Minnesota for their patience and unpublished 3D OpenMonkeyStudio pose data. We also thank Drs. Ann Kennedy, Jennifer Sun, David Anderson, and the team that created the CalMS21 data set for providing a comprehensive data set that can be used to benchmark current and future approaches to the classification of social behavior in mice. Portions of the research in this paper used the NTU RGB+D Action Recognition Dataset made available by the ROSE Lab at the Nanyang Technological University, Singapore.

## Author Contributions

JFT, AIH, MKS, and EAY wrote and reviewed the manuscript. JFT proposed the idea and AIH designed the core functionality and main analysis scripts. Both JFT and AIH created the app and worked on analysis and figures. JFT annotated the primate data. EAY and MKS provided support and funds.

## Code Availability

The app, further documentation and the open-source code written in Python can be found under (https://github.com/YttriLab/A-SOID). The code, including the code to generate these figures is open-source and available through a GitHub repository (https://github.com/YttriLab/A-SOID/tree/main/notebooks).

## Data Availability

The social mice data set (CalMS21) used in this study is available online (https://doi.org/10.22002/D1.1991) [25]. The single monkey dataset used in this study is available in our GitHub repository (https://github.com/YttriLab/A-SOID/demo dataset) thanks to the labs of Drs. Jan Zimmermann and Benjamin Hayden at the University of Minnesota. The human data set is part of the public NTU RGB+D Action Recognition Dataset made available by the ROSE Lab at the Nanyang Technological University, Singapore [39].

## Competing Interests

The authors declare no competing interests.

## Methods

### Data processing feature extraction

For the 10 selected key points, all key points but the ears from the CalMS21 resident-intruder assay, the per-frame spatiotemporal features consisted of 45 distances (**D**) and angular change (Θ) measures, and 10 total displacements (**L**) were computed. We tested alternative key point configurations in the interest of determining their effect on performance in A-SOiD. This can be of particular interest to experimenters that want to save time by including fewer body parts during pose estimation training. The performance of A-SOiD and one-shot methods were remarkably similar regardless of whether we used only the three midline key points per animal (snout, neck, tail base), five (inclusion of the hips), or seven (inclusion of the ears) per animal. These data suggest that experimenters may improve their efficiency by consolidating their key points to those that are most informative and highlight the importance of selecting the informative key points with purpose.

As described, the social behavior described in this data set can be reliably described based on a set of intra- and inter-animal features. These features are based on the distance, angular change, and speed of the animal’s body parts in relation to itself (speed, angular change) or to another body part, including its counterpart on the con-specific. For a detailed description refer to Supp. Table 1. This process is described in Algorithm 1 and the process pipeline diagram (Fig. 2a).

We then applied a temporal window with the realization that analyzing frame-to-frame differences often produces two undesirable consequences (see Fig. 2d, [9, 51]). First, with increasing sampling rates, the range of possible speeds becomes greatly reduced, e.g., a mid-stride paw may only move a few pixels with a sufficient sampling rate. Second, like any sampling modality, pose estimation comes with some degree of jitter. As such, the reported position of a stationary paw may register as moving a few pixels between each frame. Thus, to improve the signal-to-noise ratio in the CalMS21 social mice data set, we analyzed the duration distribution of these annotated behaviors with non-overlapping windows to integrate signal over, and then either sum (displacement, angular change) or averages (distance) over all samples within a given window.

We selected the 10-percentile of all bout durations to estimate a reasonable temporal resolution for the given behaviors, which resulted in a 400 ms window and 200 ms window for the CalMS21 and monkey data set, respectively. As for annotation selection, we identified the most common annotation in that time window (mode). In the rarest case of tie-breakers, we used the smaller values and did not observe any difference by using a different method. A comparison of different feature window sizes and their effects on the classifier’s performance and on the distribution of behaviors in an unsupervised embedding can be seen in Fig. 2d and Supp. Fig. 2c, respectively.

#### Algorithm 1: Feature extraction for *N* pose estimates

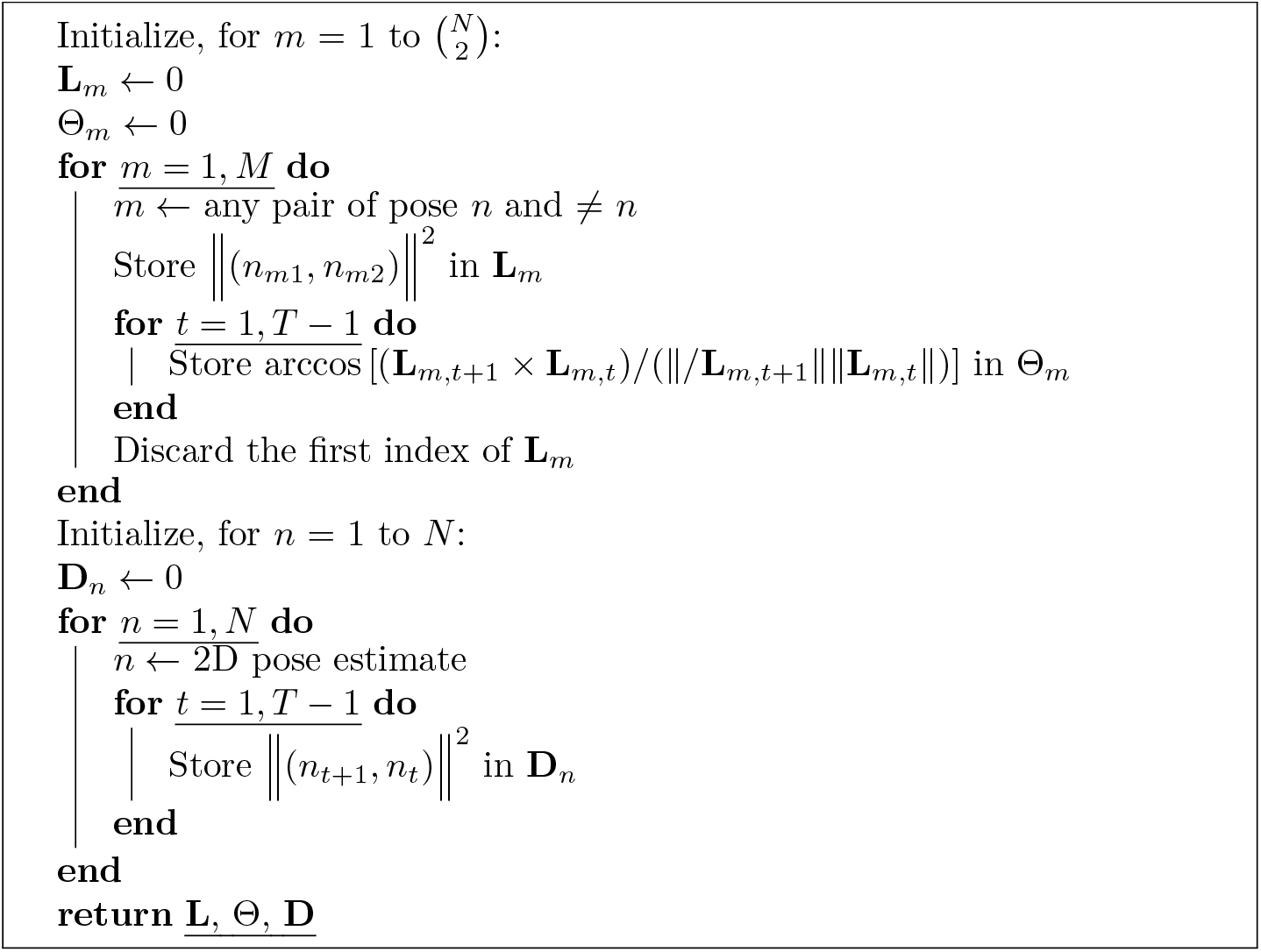

### Random forest classifier for accurate and fast prediction

To create a reproducible mapping between extracted features and aligned downsampled annotations, Random forest classifier design was chosen for high-dimensional pose relationships mapping to discrete multi-class behaviors. Random forest was iteratively implemented in the ‘Classify’ step in A-SOiD UI. Embedded in this step is a python implementation of ensemble, RandomForestClassifier(n estimators=200, random state=42, n jobs=-1, criterion=‘gini’, class weight=’balanced subsample’ from scikit-learn v.1.1.2 (https://github.com/scikit-learn/scikit-learn). In addition to active learning automatically balance training class sizes, the random forest initialized weights were dependent on the remaining diversity in class sizes (class weight=balanced subsample). In the first iteration, we subsampled 1% per annotation class to mimic the time budget. We then followed an autonomous active learning schedule to curate a selection of refined samples to incorporate into our training set.

### Autonomous iterative active learning

We first sampled 1% of the full annotated dataset. To examine the initial sample effect, we ran random under-sampling 20 times with a different seed, hence the 20 seeds described throughout the paper. Upon initializing the classifier with 1% of each annotated class, we predict the probability for all training samples. In theory, similar training samples to the initial 1% would have a high predicted probability for one class over the rest. However, if there exists a sample that does not have a predicted probability *>* 0.5 for any of the classes, we defined it as a low-confidence sample. These samples appear to be very similar in high feature dimensional space, as well as the reduced dimensional space (Fig. 1 h,i bottom). The active learning iterations were stopped when the algorithm could not extract any more confusing samples. In an iterative manner, we incorporate the supposed annotation aligned with these low-confidence samples (maximum number of samples per iteration = 200) to create a meaningful training data set. To test the model’s performance generalizability, we used the same 20% held-out test data.

### Frameshift prediction paradigm

Many end users may wish to apply the algorithm to higher frame-rate video. Because A-SOiD applies a temporal constraint depending on the temporal scale of user annotations, we designed A-SOiD to predict along a sliding window. This is mathematically implemented using offsets, pseudocode in Algorithm 2.

#### Algorithm 2 Frameshift implementation for *F* times higher sampling rate than 10fps

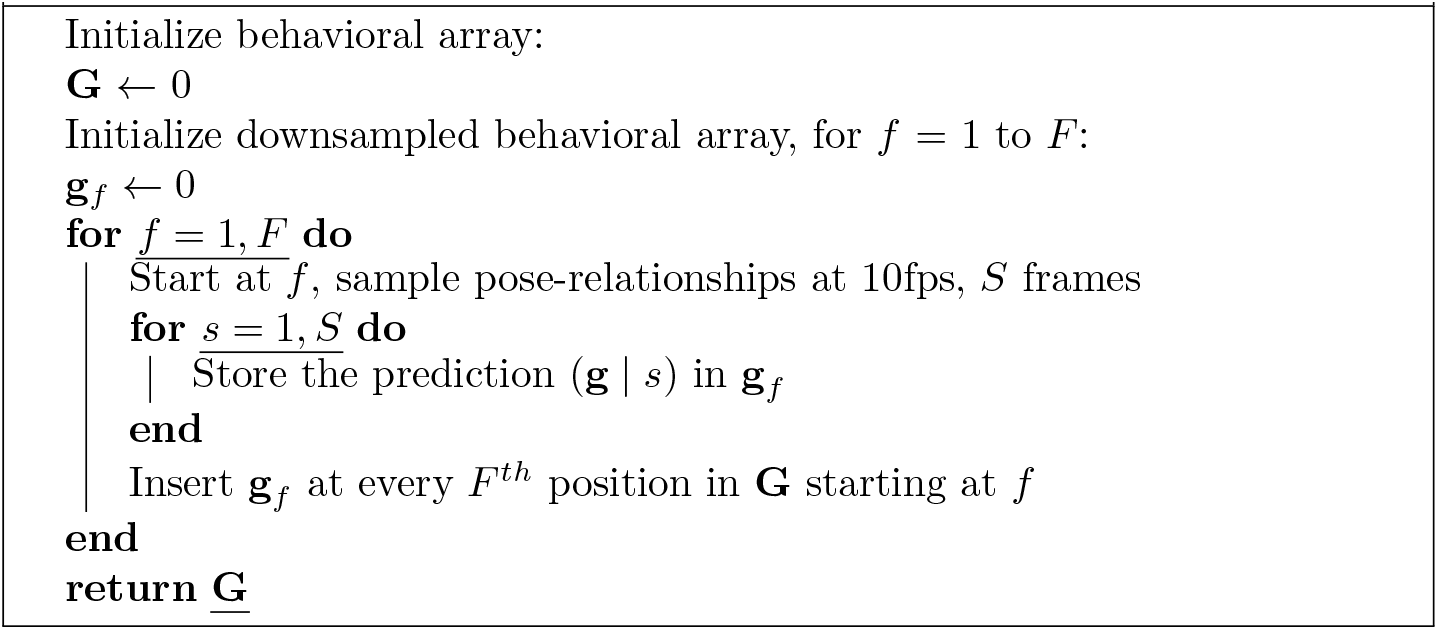

### Identify group assignments with UMAP and HDBSCAN

A-SOiD then projects the computed pose relationships (D, Θ and L) into a low-dimensional space, which facilitates behavioral identification without simplifying the data complexity. In simpler terms, a similar mouse multi-joint trajectory will retain its similarity visualized in the low-dimensional space. A-SOiD achieves this through UMAP, a state-of-the-art algorithm that utilizes Riemannian geometry to represent real-world data with the underlying assumptions of the algebraic topology [52]. UMAP, as previously described in Hsu et al. [9], is chosen over the popular t-SNE for its advantage in computational complexity, outlier distinction, and most importantly, preservation of longer-range pairwise distance relationships [52–55]. Embedded in this step is a Python implementation of umap-learn v.0.5.3 (https://github.com/lmcinnes/umap). Since our goal is to use UMAP space for clustering, we enforced the following UMAP parameters: (n neighbours=60, min dist=0.0, euclidean distance metric). In terms of n components, we call the python implementation of decomposition.PCA() from scikit-learn v.1.1.2 (https://github.com/scikit-learn/scikit-learn) and set n components to explain 0.7 of total pose estimation variance. UMAP embeddings were then clustered through the HDBSCAN algorithm [9, 56]. It is particularly useful for UMAP outlier detections as it recognizes subthreshold densities. Embedded in this step is a python implementation of hdbscan v.0.8.28 (https://github.com/scikit-learn-contrib/hdbscan). We enforced the following HDBSCAN parameters: (min cluster size=a range of 2 *−* 2.5% of the size of the data, whichever yields the most groupings).

### Extended embedding

To test different embedding techniques on the CalMS21 data set (see Supp. Fig S1 a), we used the Python implementations from scikitlearn 1.3.0 (https://scikit-learn.org) for principle component analysis (PCA) sklearn.decomposition.PCA(n components=2), as well as t-Distributed Stochastic Neighbor Embedding (t-SNE) sklearn.manifold.TSNE(n components=2).

### Motion Energy

The term “motion energy” as previously described was first introduced by Stringer et al. [41] and refers to the absolute value of the difference of consecutive frames. Since the animals are freely moving in the environment, an initial pose alignment is necessary. For this, the intruder’s neck and tail base coordinates are used. Following image registration using the estimated outline of both animals at the start of each identified behavior, we compute the motion energy (ME, absolute value of the difference of all consecutive frames) within a bout using the numpy [57] functions *np.diff*, *np.absolute*, and *np.mean* (for more information see ). We then performed averaging for each bout to reconstruct a single ME image per annotated class. In other words, each pixel in such a reconstructed ME image represents the average absolute difference between consecutive frames at a given pixel location.

### SHAP analysis

To interpret the differences in prediction from our random forest classifiers, an approach is to consolidate the Shapley value contribution of features that capture similar and correlated characteristics of the human rated classes. These consolidated values are then organized into biologically meaningful categories (inter-distance, speed, and angular change, for instance) to facilitate human inference. In our case, we obtained the Shapley values obtained from open-source Tree-SHAP package from scikit-learn v.0.41.0 (https://github.com/slundberg/shap).

### CalMS21 data set

#### Data set

While the data set consists of three parts [25]. For our purpose the first set (Task 1, Classical Classification) is the most relevant as it contains a complete training set of pose estimation sequences (70 sessions; total of 507,738 frames) including complete annotations of all frames. A separate test set of pose estimation sequences (19 sessions; total of 262,107 frames) is being used to benchmark against the challenge winner [6].

Note that at no point in time did we access the test set to improve algorithm performance.

#### Annotation Descriptions

The behavior annotation as described in Sun et al. 2021 was not altered ([25]). Please refer to the original publication for detailed information. Notably, the majority of annotations did not include one of the three behaviors. These widely-divergent samples were collectively annotated as “other” and are therefore not considered in evaluations regarding this benchmark data set.

#### Pose estimation

The provided pose estimation of the data set was extracted using the MARS [6]. MARS identifies seven user-defined body parts (snout, ears, neck, hips, and tail base). For more information refer to Sun et al. 2021 [6, 25]. During feature engineering, we discarded the left and right ear key points as they did not provide additional information about the underlying behavior.

#### Videos

The data set also contains 70 videos of mice resident-intruder assay, of which matches the frame-wise pose estimation and annotation of several behaviors. These videos were used to evaluate the performance of DeepEthogram.

### Non-human primate data set

A single housed monkey exploring the environment for 5 minutes. Monkey’s pose was generated by OpenMonkeyStudio [7] in line with the data in Maisson et al. 2022 [58] and Voloh et al. 2022 [**Voloh2022PrefrontalMacaques**]. All research and animal care procedures were conducted in accordance with the University of Minnesota Institutional Animal Care and Use Committee approval and in accordance with National Institutes of Health standards for the care and use of non-human primates.

#### Annotation Descriptions

Behavior annotation was done using BORIS [59] by an expert annotator based on an unpublished video displaying the 3D pose data. The animal’s behavior was separated into distinct categories that were exclusive to one another - i.e., only a single behavior could be shown at the same time.

1. **Walk**: The monkey moved across the arena using its feet or feet and hands touching the ground (floor or platform).
2. **Jump**: The monkey jumped from the ground onto a platform or from one platform to another, leaving the ground and remaining for a certain duration in the air. This included the moment preparing the “jump” and immediately after landing.
3. **Climb sidewall**: The monkey left the ground completely and moved on a sidewall of the arena using its hands and feet. This does not include moments where the monkey transfers from the ground to the sidewall to separate the behavior from rearing.
4. **Climb ceiling**: The monkey left the ground or sidewall completely and climbed on the ceiling of the arena using its hands or hands and feet. This includes moments when the monkey transfers from the sidewall to the ceiling and vice versa.
5. **Rearing**: The monkey touches the sidewall at any point while remaining on the ground or on a platform on its feet. This includes initiating and finishing the rearing until the next behavior is identified.

### Human data set

#### Data set

The NTU-RGB-D60 data set [39] contains examples for 60 different actions, including 11 social activities between two people. While the data set contains movement estimates from multiple sources, we exclusively relied on pose estimation data. The pose estimation data contains 25 key points per skeleton across the entire body of the subjects with focus on joints, hands, feet, and the spine. Note that unlike the CALMS21 and monkey data sets, this data set is highly balanced and contains an equal number of examples per action. We split the data set into single human (49 actions) and social human (11 actions) and tested our approach on them separately. The data set contains over 56k sessions (n = 56578), separated by action, over multiple subjects. If sessions marked as “single” contained more than one skeleton, we selected the first and excluded all others. If sessions marked as “social” did not contain two skeletons, we excluded the session entirely (n total = 10347, n selected = 9892). The behavior annotation as described in Shahroudy et al. was not altered, but split into 2 separate groups (single and social; see Table 1 and 2). For this, we renamed behaviors that were previously termed A50-A60, to S01-S11, but kept all other information unchanged.

**Table 1:**
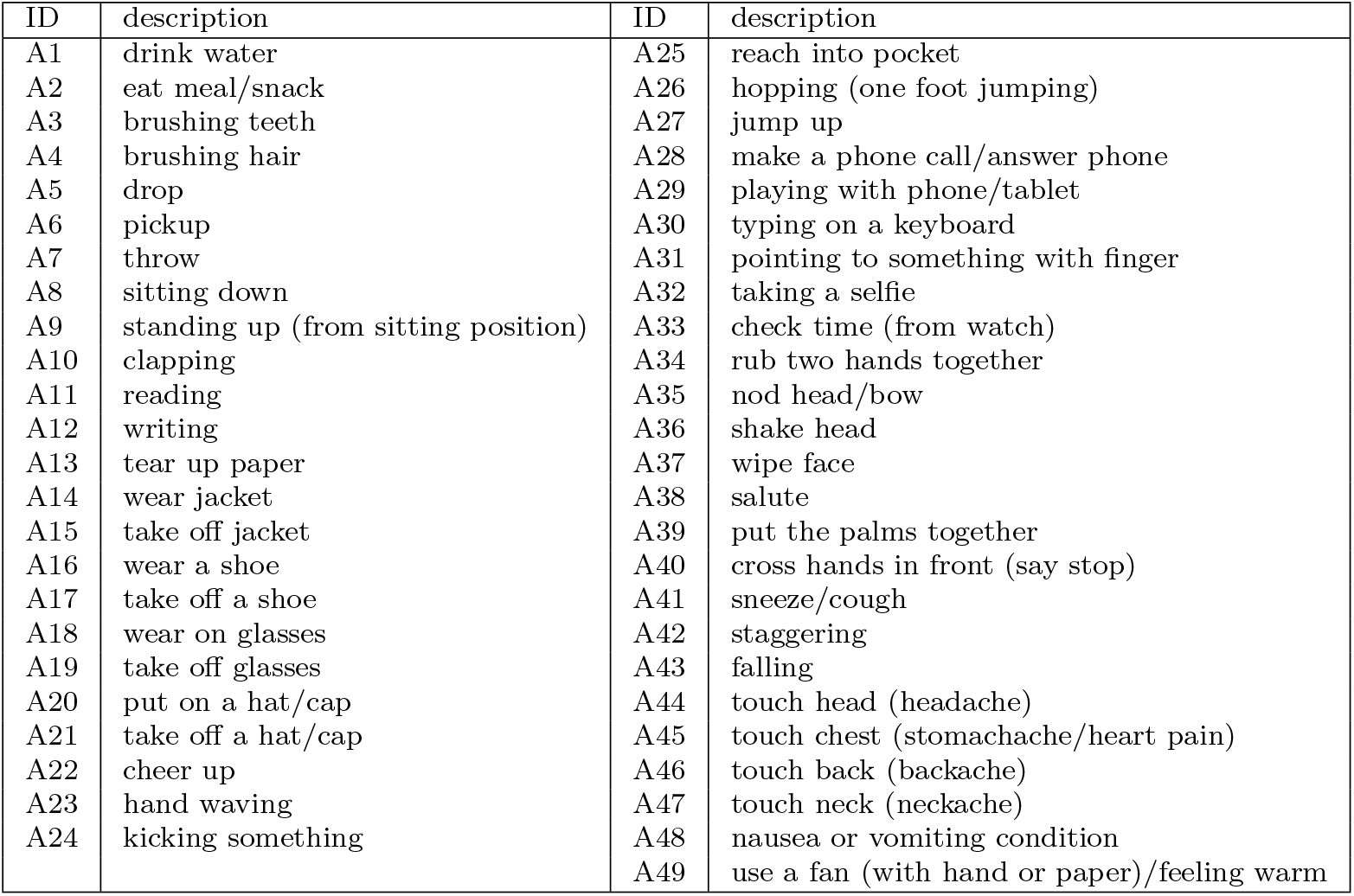
Actions included in the human data set (single)

| ID | description | ID | description |
| --- | --- | --- | --- |
| A1 | drink water | A25 | reach into pocket |
| A2 | eat meal/snack | A26 | hopping (one foot jumping) |
| A3 | brushing teeth | A27 | jump up |
| A4 | brushing hair | A28 | make a phone call/answer phone |
| A5 | drop | A29 | playing with phone/tablet |
| A6 | pickup | A30 | typing on a keyboard |
| A7 | throw | A31 | pointing to something with finger |
| A8 | sitting down | A32 | taking a selfie |
| A9 | standing up (from sitting position) | A33 | check time (from watch) |
| A10 | clapping | A34 | rub two hands together |
| A11 | reading | A35 | nod head/bow |
| A12 | writing | A36 | shake head |
| A13 | tear up paper | A37 | wipe face |
| A14 | wear jacket | A38 | salute |
| A15 | take off jacket | A39 | put the palms together |
| A16 | wear a shoe | A40 | cross hands in front (say stop) |
| A17 | take off a shoe | A41 | sneeze/cough |
| A18 | wear on glasses | A42 | staggering |
| A19 | take off glasses | A43 | falling |
| A20 | put on a hat/cap | A44 | touch head (headache) |
| A21 | take off a hat/cap | A45 | touch chest (stomachache/heart pain) |
| A22 | cheer up | A46 | touch back (backache) |
| A23 | hand waving | A47 | touch neck (neckache) |
| A24 | kicking something | A48 | nausea or vomiting condition |
|  |  | A49 | use a fan (with hand or paper)/feeling warm |

#### Feature extraction

As previously reported, we used our 3D feature extraction to extract spatio-temporal features between all body parts and skeletons. We selected a time window of 50 frames for both “single” and “social” actions after investigating the distribution of session lengths. This resulted in 43259 samples with 625 features (single) and 131277 samples with 2500 features (social).

#### Active learning

To test the generalizability of our model on held-out test data, we split the subselected data into a training and test set (80/20 split) using the python implementation of sklearn.model selection.train test split().

As previously reported, we used our active learning regime to train a classifier on low-confidence examples across multiple iterations. We initialized the classifier with 1% of training data and set the maximum number of samples per iteration to roughly 1% of available training data; 1600 (social) and 6000 (single), respectively. Further, we cross validated our active learning approach by shuffling the training data (n = 3). We then ran active learning until the classifiers performance reached the one-shot trained classifier or the majority of training data was used (social: 50 iterations; single: 90 iterations).

### Benchmarking

#### SiMBA

To evaluate SimBA’s performance on the CalMS21 data set, we utilized the pose estimation data provided. Using the provided GUI (SimBA v0.91.8) [8], we first created a custom body part model to allow the import of the data set. For this, we first converted the original pose representation of both mice (e.g., mouse 1: nose, mouse 2: nose) into a flattened representation (e.g., nose 1, nose 2) which is then imported into SimBA as two separate individuals. We used the same training and test set (70 videos:19 videos) to evaluate the classifier’s performance after training. In contrast to A-SOiD which uses a single multi-class classifier, SimBA employs a number of binary classifiers (one for each behavior). Notably, SimBA expanded their set of available key point configurations and optional features since their initial release. These additions allow advanced users to fine-tune their model parameters, which in turn can lead to increased performance levels if done correctly. While these optional features include over- and undersampling techniques as well, we opted to compare SiMBA’s performance with a set of standard parameters in order to allow a direct comparison between the two solutions. To train the classifiers (RandomForest) for each behavior, we used a set of standard parameters (train test size = 0.01, under sample setting = None, over sample setting = None, rf n estimators = 200, rf min sample leaf = 1, rf max features = sqrt, rf n jobs = -1, rf criterion = gini) across all instances. After training, we used the test set to predict annotations, of which we took the max prediction probability across the 4 binary classifiers, and evaluated the performance the same way as the other solutions.

#### DeepEthogram

To evaluate the performance of Deepethogram’s (DEG) performance on the CalMS21 data set, we utilized the video material provided. We split the videos into the same training and test set as used in the remaining benchmark (70 videos:19 videos). Following the DEG workflow, we used the training set to train the three DEG models using their provided UI (v0.1.4) [33]. Using Pytorch 1.13 and Cuda 11.7, the models (FlowGenerator: 200221 115158 TinyMotionNet, FeatureExtractor: 200415 125824 hidden two stream kinetics degf, and Sequence) were trained with default parameters on a workstation (Titan RTX 21 Gb GPU; 384 Gb RAM; Intel Xeon CPU). After training, we used the test set to predict labels for each given video. Given DEG’s capabilities to predict multiple behaviors within the same frame - i.e., behaviors are not exclusive in their workflow, and DEG does not export prediction probabilities; extracting a single label in instances with multiple predictions was required. In these edge cases, we selected labels randomly from the set of predictions. We then used the resulting labels to evaluate the performance in the same way as the other solutions. Note that in DEG a background class is preassigned so that we set all “other” annotations to said “background” in order to align both methods.

### Performance evaluation

With regards to A-SOiD, we optimized our features based on previous experience with B-SOiD on single animals. We did not optimize performance based on test set performance.

To evaluate the performance of each model, we calculated the unweighted F1 score for each behavior, excluding “other” in the case of the CalMS21 data set as previously established. We used the Python implementation of scikit-learn 1.3.0 (https://scikit-learn.org) sklearn.metrics.f1 score(). To calculate the average F1 score across classes (e.g., during comparison between different approaches), we calculated the unweighted mean of individual F1 scores.

To calculate the confusion matrix, we used the Python implementation of scikit-learn 1.3.0 (https://scikit-learn.org) sklearn.metrics.confusion matrix() .

### Runtime evaluation

Given that A-SOiD decreases the number of training samples required via the selection of low-confidence examples and that different datasets consist of unique combinations of body parts (effecting feature dimensions) and sample distributions for user-defined behaviors, the algorithm’s runtime is a complex interplay of many factors. Therefore, we estimate the runtime performance using subsets of the benchmark dataset (CalMS21) and extrapolate the runtime with increasing sample sizes beyond the original data set given the same features to give users a principle idea of A-SOiD’s runtime performance.

We ran the core A-SOiD scripts, responsible for feature extraction, active learning, and evaluation in a Google Collab notebook (Free version, 12.7 Gb RAM) to best control for variations in background workload.

## Notes

https://github.com/YttriLab/A-SOID

### Competing Interest Statement

The authors have declared no competing interest.

### Summary of Updates

In the revised version we benchmarked our approach against standard methods in the field. In addition, we included a new dataset including the classification of social interactions in human subjects.

