## supplementary data for "A-SOiD, an active learning platform for expert-guided, data efficient discovery of behavior"

### Extended discussion

#### Data complexity can be resolved by active learning

In studies investigating the behavior of animals in social contexts, approaches that have been trying to disentangle the high data complexity of multiple interacting individuals (unsupervised), oftentimes struggle to produce results that align with human consensus. This is partly because many existing solutions are predisposed to focus on single animal behaviors [**Luxem2022IdentifyingMotionb**, 1–5] as the within-animal spatiotemporal dynamics are generally more conserved and less complex than intra-animal dynamics.
On the other hand, most supervised solutions require a sizable ground truth data set to reliably reproduce human expectations. One reason for this is that most collected data sets are inherently unbalanced. They consist of high-frequency behaviors (e.g., investigation) accompanied by low-frequency behaviors (e.g., attack). Behaviors that are underrepresented in the data set are then underrepresented in training and will result in poor performance levels. In both cases, therefore, the complexity of the expected outcome and the composition of the data inhibit the reproduction of the human reference frame.
A-SOiD solves this challenge by employing an active learning regime. By explicitly refining low-confidence predictions, the algorithm focuses on unclear decision boundaries between classes and continuously learns to reproduce the expert's definitions with high consistency (Fig. 1h-i, bottom). By starting with an absolute minimum number of annotations (yet still maintaining the size differences), we let A-SOiD determine which features are to be annotated. This approach effectively sparsifies data means while focusing on the outliers. Over iterations, the training data is therefore auto-balanced across classes. This capacity eradicates the need for additional data augmentation or filtering steps to balance the underlying data (see Supp. Table 2, Fig. 3 d-f). Additionally, this approach reduces necessary ground truth annotation for uncommon, low-frequency behaviors (Fig. 2c) and the cost required to implement such a solution in a dynamic analysis pipeline.
However, a general challenge of behavioral analysis is the temporal scale at which behavior forms and changes in observed animals. Specifically, behavioral expressions can be subject to changes through previous experiences and different contexts. The possible range of observable behavior is further limited by the context and duration of each recording session. Therefore, it is unrealistic for most projects to collect an extensive ground truth encompassing the entire behavioral repertoire. Insufficient ground truth data may lead to inaccurate predictions at different times in the animal's life or across a wide range of behavioral assays. Here, an optimal solution would be able to continuously build upon the observed repertoire and retrain the algorithm with novel variations. Unfortunately, many approaches, especially unsupervised, would require a complete restart and realignment to human reference frames, increasing the cost to employ such methods drastically. In contrast, iterative approaches such as A-SOiD have the potential to be continuously updated

with new observations using the active learning scheme.

#### Utility of creating an equally emphasized network

Further explanation of a behavioral classifier can provide insight into differences in predictive performance, which subsequently can be used to understand behavioral differences between experimental conditions. A classifier can predict at a similar overall performance with a wide variety of training regimes. It is then required to dissect the model independently. Recent work on explainable AI [6, 7] has allowed machine learning engineers to rank features given the labels. In addition to better performing small classes, the benefit of having a balanced representation is for explanatory approaches like SHAP to independently identify behavioral differences without being subject to overprioritizing one class over the other. In our results, we have demonstrated albeit similar performing models, our iterative learning schemes provided a precise separation between feature value and its impact on the model (SHAP value). If we used an imbalanced dataset to fit a classifier, the feature value impact on model output will be intermixed since all the original model had to do was overclassifying this large class.
Strategies to increase the transparency of behavioral classification to quickly assess the differences between discovered patterns (e.g., behavioral sub-types), we previously employed motion energy (see Methods) [1, 8]. Motion energy is an intuitive and informative way to generate a visual summary of the action within a found cluster (Fig. 3b-c). In our hands, we utilized motion energy images to quickly differentiate sub-types of anogenital investigation (Fig. 3). Note that further analysis can also be done by comparing variability within and across groups providing a valuable statistic for cluster quality [1]. Another approach is using SHAP-based reporting of the underlying feature importance, which can help to share and compare conserved patterns across studies (for a review, see [9]). In this study, we employed SHAP-analysis to investigate the learning process across multiple iterations during active learning and could identify that specific sets of features accounted for the increased performance of our classifiers (compare Rank 1-10 features across iterations in Fig. 2e-h and Suppl. Fig. S4).

### Active learning framework for user-defined data

To allow the integration of our developed approach into already existing behavior analysis pipelines, we created a streamlit-based application that integrates the core features of A-SOiD into a user-friendly, no-coding required GUI solution that can be downloaded and used on custom data. For this, we developed a multi-step pipeline that guides users, independently of their previous machine-learning knowledge, through the process of generating a well-trained, semi-supervised classifier for their own use case. While the underlying code is based on the open-source language Python and available on GitHub (https://github.com/YttriLab/A-SOID), the use of A-SOiD's core feature

(active learning) to reduce the amount of necessary ground truth data considerably can be directly used by installing the app on a local computer. In general, users are required to provide a small labeled data set (ground truth) with behavioral categories of their choice using one of the many available labeling tools (e.g., BORIS; [10]) or import their previous supervised machine learning data sets. Following the upload of data (see Supp. Fig. S1a), an A-SOiD project is created, including several parameters that further enable users to select individual animals (in social data) and exclude body parts from the feature extraction. Based on the configuration, the feature extraction (Supp. Fig. S1 c) can be further customized by defining a "bout length" referring to the temporal resolution in which single motifs are expected to appear (e.g., the shortest duration of a definable component of the designated behavior is expected to last; see also Fig. 1). The extracted features are then used in combination with the labeled ground truth to train a baseline model. Here, an initial evaluation will give users insight into the performance of their base data set (Supp. Fig. S1 d). Note that different splits are used to allow for a more thorough analysis (see Methods for further details).
The first iteration (iter0) will be trained on a subset of the available training data and is then used as a basis for the following active learning iterations (automatic active learning), A-SOiD automatically selects new samples from the remaining training set for each iteration until no low-confidence examples are left, or the maximum number of iterations has been reached.
Finally, users can upload and classify new data using the app and the previously trained classifier (Supp. Fig. S1e). To gain further insight into the results of the classification, the app offers a reporting tab that allows users to view and export a selected set of analysis reports, including the common ethogram and statistics (Supp. Fig. S1f).

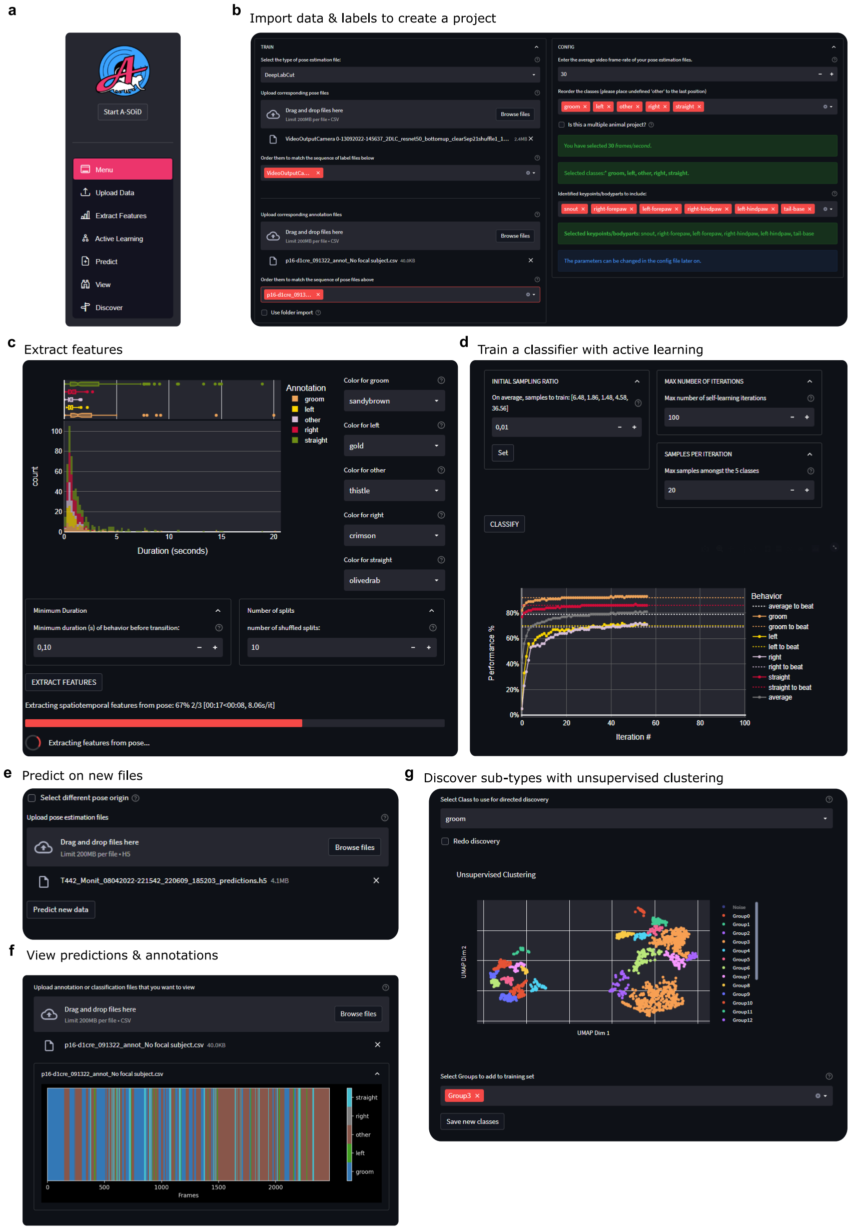
a
Start A-SOiD
Menu
Upload Data
Extract Features
Active Learning
Predict
View
Discover
b
Import data & labels to create a project
c
Extract features
Annotation
groom
left
other
right
straight
count
Duration (seconds)
Color for groom
sandybrown
Color for left
gold
Color for other
thistle
Color for right
crimson
Color for straight
olivedrab
Minimum Duration
Minimum duration (s) of behavior before transition:
Number of splits
number of shuffled splits:
EXTRACT FEATURES
Extracting features from pose...
d
Train a classifier with active learning
INITIAL SAMPLING RATIO
MAX NUMBER OF ITERATIONS
Max number of self-learning iterations
SAMPLES PER ITERATION
Max samples amongst the 5 classes
Set
CLASSIFY
Behavior
average to beat
groom
groom to beat
left
left to beat
right
right to beat
straight
straight to beat
average
Performance %
Iteration #
e
Predict on new files
Select different pose origin
Upload pose estimation files
Drag and drop files here
Limit 200MB per file • H5
Browse files
T442_Monit_08042022-221542_220609_185203_predictions.h5
Predict new data
f
View predictions & annotations
Upload annotation or classification files that you want to view
Drag and drop files here
Limit 200MB per file • CSV
Browse files
p16-d1cre_091322_annot_No focal subject.csv
Frames
g
Discover sub-types with unsupervised clustering
Select Class to use for directed discovery
groom
Redo discovery
Unsupervised Clustering
UMAP Dim 1
UMAP Dim 2
Select Groups to add to training set
Group3
Save new classes

Figure S1: Active learning framework on user-defined data. a) The A-SOiD GUI offers step-by-step navigation to run A-SOiD on your own data. b) First users can select the origin/type of their pose estimation data (SLEAP, DLC, or CALMS21) and uploads their data set including a previously labeled ground truth. Using a user-defined working directory and prefix, previous sessions can be continued at later stages by uploading the corresponding config file (not shown). Right, the user can enter basic parameters (framerate, resolution), behavioral categories of interest that are contained in the ground truth data set as well as sub-select individuals/animals and key points/body parts as a basis for feature extraction. c) After input of a temporal reference frame (aka. bout length) for feature extraction using a histogram as shown in Fig. 1 (top), features are extracted, and a number of splits are provided to evaluate later classification training. d) In the active learning segment, a classifier is trained as described previously by the iterative addition of low-confidence predictions. Here, refinement is directly taken from the remaining ground truth. During each iteration, the model's performance is evaluated on a held-out test data for multiple splits. This process can be viewed live for each iteration. e) Finally, once the training is complete, users can use the app to upload new unlabeled data and use the previously trained model for classification. f) After classification, the app allows users to go through the results and view a brief report. g) Users are also able to discover conserved patterns in their ground truth data by selectively clustering annotation classes with directed unsupervised classification. Sub-types of interest can then be exported to create a new training set and be used to train a classifier.

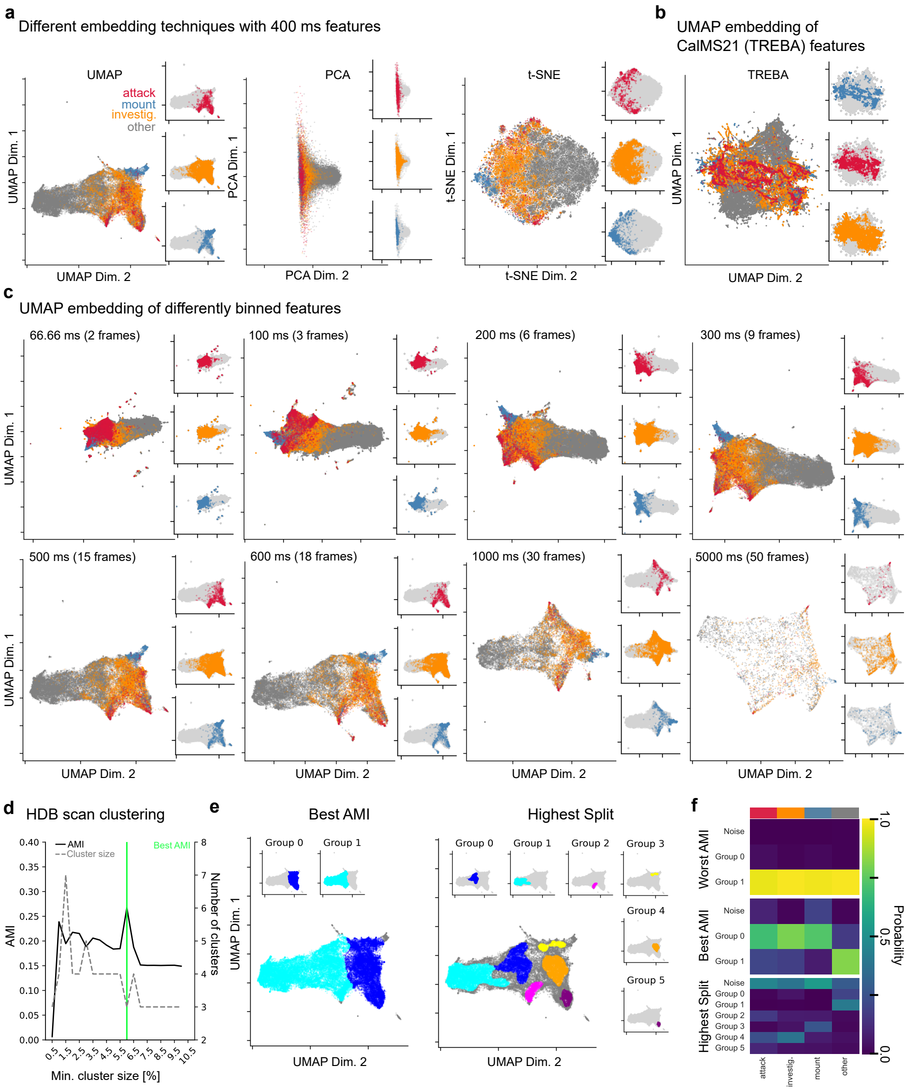
a
Different embedding techniques with 400 ms features
UMAP
attack
mount
investig.
other
UMAP Dim. 1
UMAP Dim. 2
PCA
PCA Dim. 1
PCA Dim. 2
t-SNE
t-SNE Dim. 1
t-SNE Dim. 2
b
UMAP embedding of
CalMS21 (TREBA) features
TREBA
c
UMAP embedding of differently binned features
66.66 ms (2 frames)
100 ms (3 frames)
200 ms (6 frames)
300 ms (9 frames)
500 ms (15 frames)
600 ms (18 frames)
1000 ms (30 frames)
5000 ms (50 frames)
d
HDB scan clustering
AMI
Cluster size
Best AMI
Number of clusters
Min. cluster size [%]
e
Best AMI
Highest Split
Group 0
Group 1
Group 2
Group 3
Group 4
Group 5
f
Worst AMI
Best AMI
Highest Split
Noise
Probability
attack
investig.
mount
other

Figure S2: CalMS21 Task 1 extended embeddings and feature binning. a) Different embeddings of the features extracted from the CalMS21 data set used in our active learning approach (feature window = 400 ms or 12 frames). Left, original 2D UMAP embedding as seen in Figure 1g. Middle, principal component analysis (PCA, n = 2). Right, t-Distributed Stochastic Neighbor Embedding (t-SNE, n = 2). The annotations for each underlying behavior are superimposed onto the embedding (attack = red, investigation = orange, mount = blue, other = dark grey). Inserts show the underlying distribution separated by behavior to allow visual inspection of overlapping samples in the total embedding (light grey). b) 2D UMAP embedding of the original CalMS21 Task1 features provided with the data set. The features for the baseline model include the pose estimation of all body parts and 32 additional features extracted with TREBA (Sun et al. 2020). c) To examine the effect of differently sized feature bins, we plotted the 2D UMAP embeddings across a range of 2 frames to 150 frames. Note that 66.66 ms or 2 frames (top left) is the minimum amount of temporal binning that can be achieved with a framerate of 30 Hz, and a temporal binning of 5 sec or 150 frames (bottom right) is considered very coarse for most observations. d) Using a hyperparameter search strategy, we calculated the adjusted mutual information score (AMI) for a minimum cluster size range of 0.5 to 10.0% with 0.5% intervals. For this, we calculated the AMI score (black line) between the assignments and the original human annotations for each step to assess whether the clusters reproduce the human reference frame of the underlying data. An AMI score of 1.0 indicates a perfect overlap of mutual information even if the number of groups is different- i.e., if investigation was perfectly represented by two clusters instead of one, the mutual information would still be 1. In addition, we looked at the total number of clusters, including noise (grey, dashed line) resulting in each step, as we expected a higher split to be more likely to incorporate the subtle differences between the behaviors. e) Selected HDBscan clustering of the embedded features (400 ms) in d. Left, the highest AMI reached a minimum cluster size of 6.0% (AMI = 0.267). Right, the highest number of clusters was found with a minimum cluster size of 1.5% (AMI = 0.195). Colors show identified clusters within each individual plot. Note that samples that cannot be confidently associated with a cluster are collectively annotated as noise (dark grey) and can therefore span the entire embedding. f) 2D histogram of the assigned cluster groups (y-axis) in d in relation to their ground truth annotations. Each histogram is normalized so that the sum of each column (ground truth behavior, e.g., attack) is 1.0.

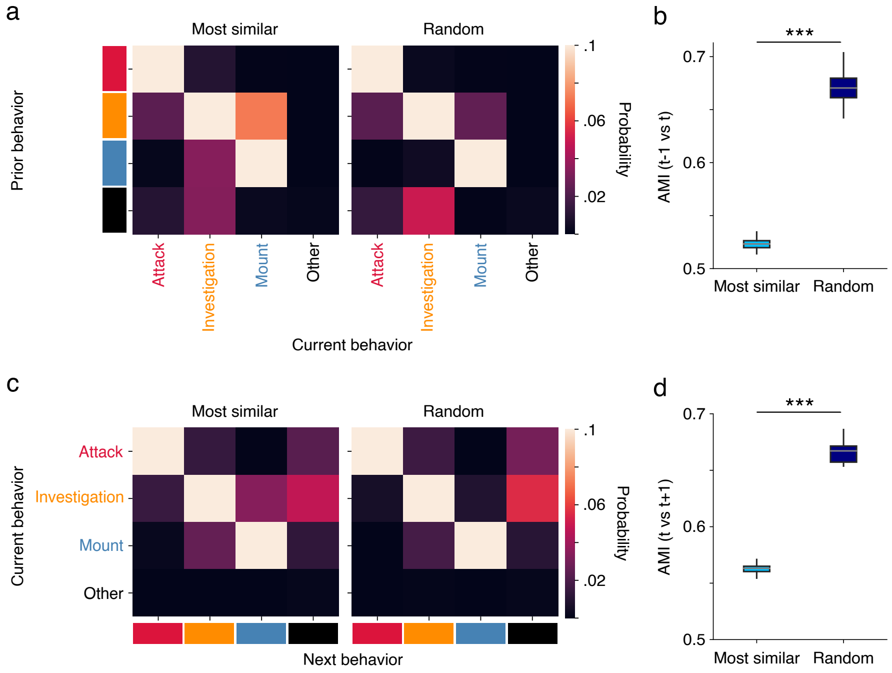


Figure S3: More candidates for active learning refinement appear at behavior transitions. a) Transition matrix for the frame annotation that happens before refinement candidates throughout A-SOiD (left), when compared to a random frame selection of these behaviors (right). b) Adjusted mutual information score as a metric to quantify similarity between prior frame (t-1) and refinement candidate/random selection (t). c) Transition matrix for the frame annotation that happens after refinement candidates throughout A-SOiD (left), in contrast to a random frame selection of these behaviors (right). d) Adjusted mutual information score as a metric to quantify similarity between next frame (t+1) and refinement candidate/random selection. Wilcoxon rank sum test, *** = $p < 0.001$.

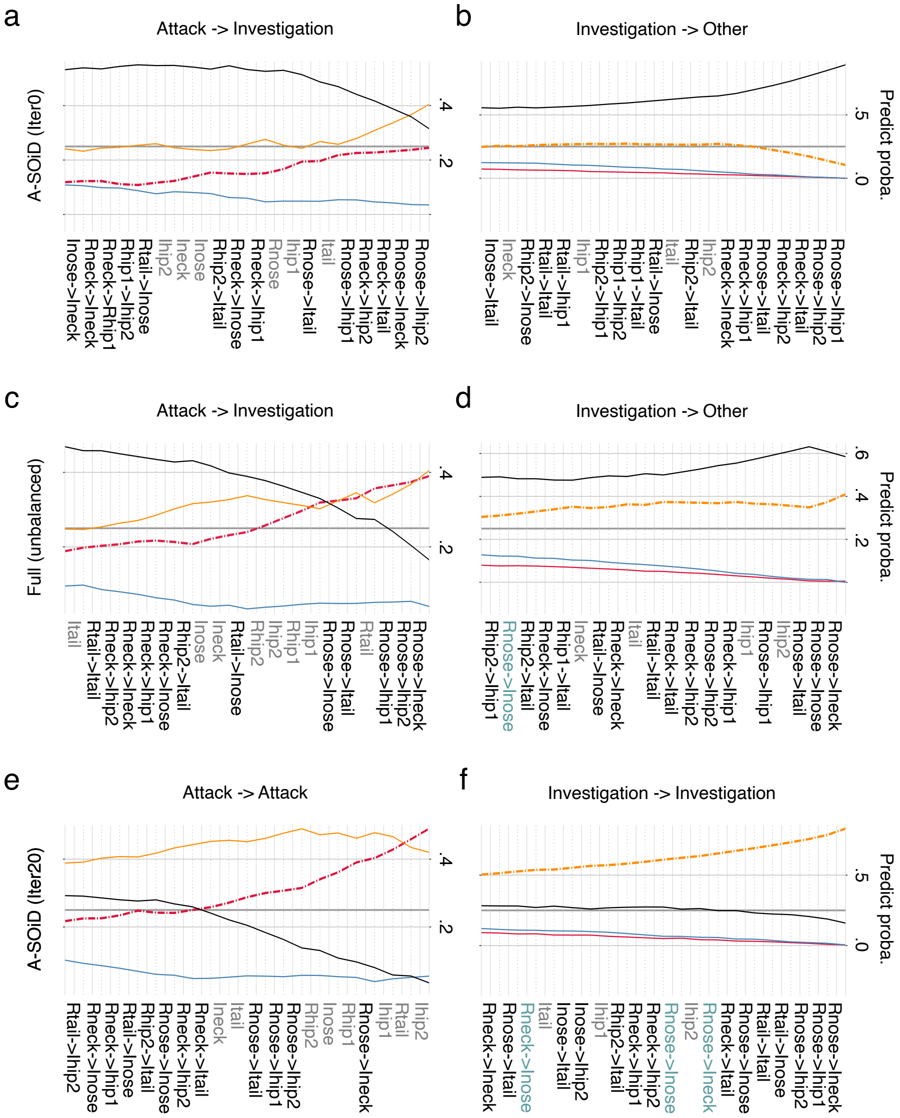


Figure S4: SHAP-replicated decision paths of two instances where A-SOiD, but not using full training data at once, correctly predicted "attack" and "investigation". a) Multi-output decision plot on an instance where human annotated "attack", but A-SOiD iteration 0 predicted "investigation". The features on the x-axis are ranked sorted by their impact on cumulative prediction probability on the y-axis. The highest final prediction probability of the four color-coded lines revealed the prediction was "investigation".

Figure S4: b) Multi-output decision plot on an instance where human annotated "investigation", but A-SOiD iteration 0 predicted "other". The features on the x-axis are ranked and sorted by their impact on cumulative prediction probability on the y-axis. The highest final prediction probability of the four color-coded lines revealed the prediction was "other". c) Similar to a), a multi-output decision plot on an instance where human annotated "attack", but full training data at once incorrectly predicted "investigation". d) Similar to b), a multi-output decision plot on an instance where human annotated "investigation", but full training at once incorrectly predicted "other". e) Similar to a) and c), a multi-output decision plot on an instance where human annotated "attack", and A-SOiD iteration 20 correctly predicted "attack". f) Similar to b) and d), a multi-output decision plot on an instance where human annotated "investigation", and A-SOiD iteration 20 correctly predicted "investigation". Red: "attack"; Orange: "investigation"; Blue: "mount"; Black: "other". Features names are black when they're inter-distance, gray when they're speed, and teal when they're angular change. R: resident; I: intruder.

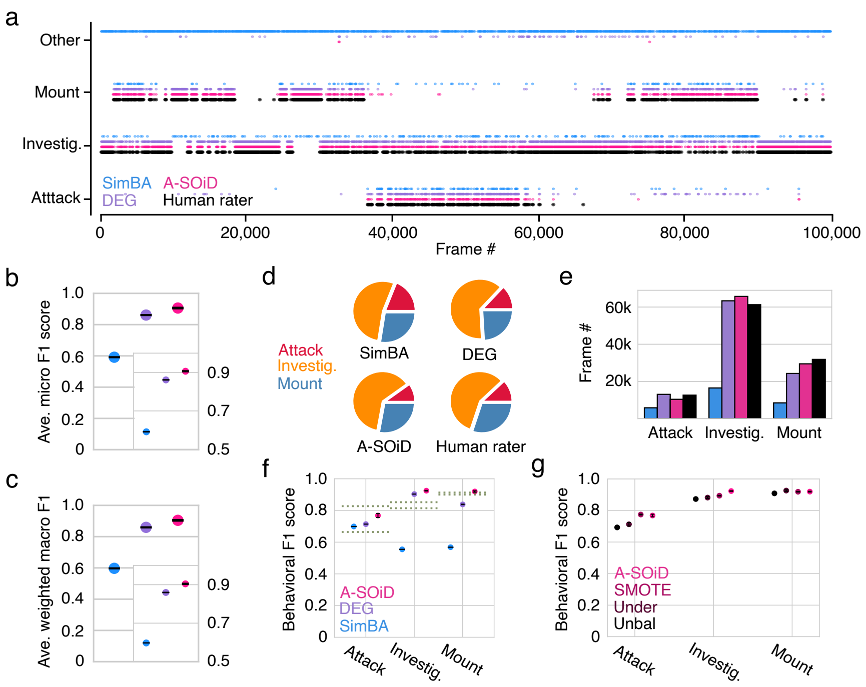


Figure S5: Benchmark results against state-of-the-art supervised classification methods. a) Predicted ethogram (including "other") using SimBA (blue), DeepEthogram (DEG, purple), and A-SOiD (pink) against target human annotation (black) without "other". Every fiftieth frame is shown to allow for better visualization. Session n=19. b) Micro F1 scores across all behaviors (including classification of "other") using SimBA (blue), DEG (purple), and A-SOiD (pink). Error bars represent */pm* 3 SD across 20 seeds. c) Weighted macro F1 scores averaged across all behaviors (including classification of "other") using SimBA (blue), DEG (purple), and A-SOiD (pink). Error bars represent */pm* 3 SD across 20 seeds. d) The percentages of each behavior predicted for each algorithm (top left: SimBA; DEG: top right; A-SOiD: bottom left; human annotation: bottom right). e) The total number of frames being predicted as "attack", "investigation", "mount", and "other", for each algorithm (top left: SimBA; DEG: top right; A-SOiD: bottom left; human annotation: bottom right). f) F1 scores for individual behavior using SimBA (blue), DEG (purple), and A-SOiD (pink). Error bars represent */pm* 3 SD across 20 seeds. g) F1 scores for individual behavior using unbalanced (unbal, black), random under-sampling (under, brown), SMOTE (maroon), and A-SOiD (pink). Error bars represent */pm* 3 SD across 20 seeds.

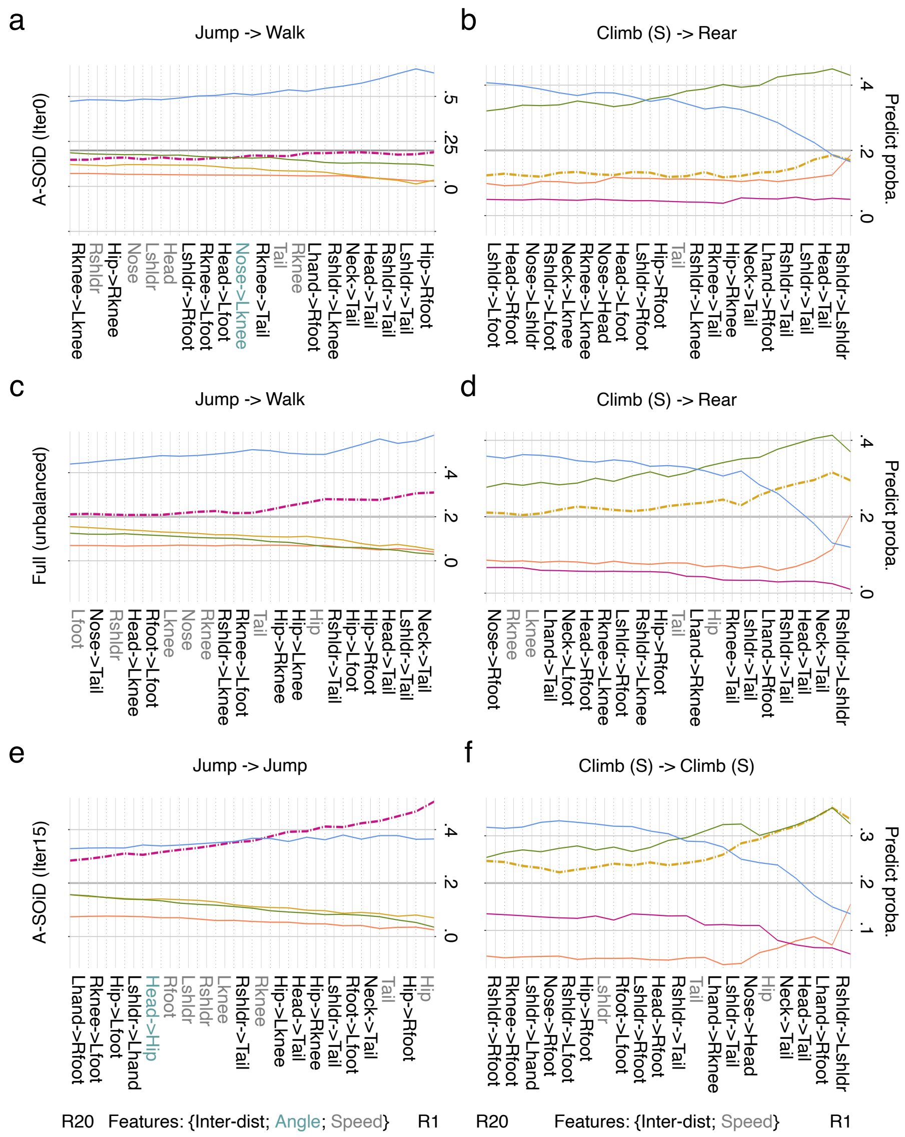

Figure S6: SHAP-replicated decision paths of two instances where A-SOiD, but not using full training data at once, correctly predicted "jump" and "climb (S)". a) Multi-output decision plot on an instance where human annotated "jump", but A-SOiD iteration 0 predicted "walk". The features on the x-axis are ranked and sorted by their impact on cumulative prediction probability on the y-axis. The highest final prediction probability of the four color-coded lines revealed the prediction was "walk".

Figure S6: b) Multi-output decision plot on an instance where human annotated “climb (S)”, but A-SOiD iteration 0 predicted “rear”. The features on the x-axis are ranked and sorted by their impact on cumulative prediction probability on the y-axis. The highest final prediction probability of the four color-coded lines revealed the prediction was “rear”. c) Similar to a), a multi-output decision plot on an instance where human annotated “jump”, but full training data at once incorrectly predicted “walk”. d) Similar to b), a multi-output decision plot on an instance where human annotated “climb (S)”, but full training at once incorrectly predicted “rear”. e) Similar to a) and c), a multi-output decision plot on an instance where human annotated “jump”, and A-SOiD iteration 20 correctly predicted “jump”. f) Similar to b) and d), a multi-output decision plot on an instance where human annotated “climb (S)”, and A-SOiD iteration 20 correctly predicted “climb (S)”. Orange: “ceiling climb” (Climb C); Yellow: “sidewall climb” (Climb S); Pink: “jump”; Green: “rear”; Blue: “walk”. Features names are black when they’re inter-distance, gray when they’re speed, and teal when they’re angular change. R: right-sided body parts; L: left-sided body parts.

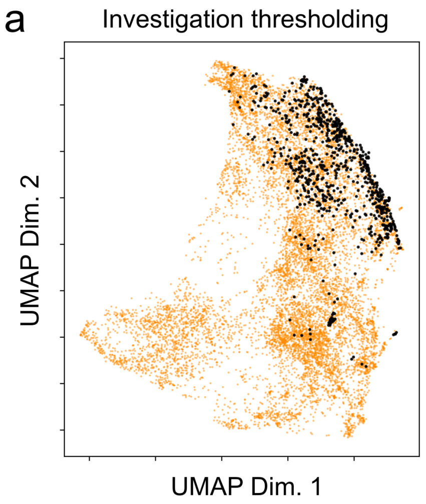


Figure S7: Unsupervised embedding of the investigation class can be explored using heuristic approach a) Annotation of all data points within the investigation class in which the distance between the resident’s snout and the intruder’s tail base was lower than a manually set threshold (15 pixels ). A comparison revealed an extensive overlap between the unsupervised clusters of sub-class 2 and sub-class 5 overlap with the top-down, manually selected feature space (compare with Fig. 3a).

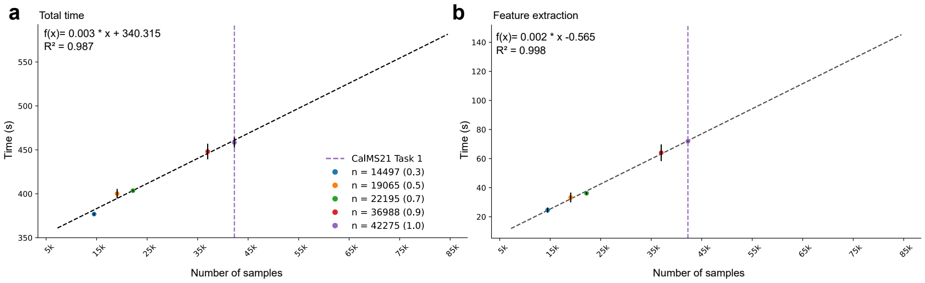


Figure S8: Active learning speed across different sample sizes. To estimate the total time it takes to run larger datasets with A-SOiD, we timed the time it takes for our active learning regime (max iterations = 20, max number of samples per iteration = 200, initial ratio = 0.01) across a range of subsets (0.3 to 1.0) of the CalMS21 data set (number of features = 100). We then fit a linear function to the measurements to estimate the performance speed with increasing sample sizes. a) Total time A-SOiD takes to complete 20 iterations, including feature extraction of the train set. Given the fit, every 1000 new samples increase the runtime by 3 seconds. The time it takes to run 1 Million samples is roughly 53 min. b) Isolated feature extraction speed for each subset. Given the fit, every 1000 samples increase the runtime by 2 seconds ( 1M samples about 28 min). Notably, we considerably optimized feature extraction by employing just-in-time compilation using the Python implementations of numba 0.52.0 (https://github.com/numba/numba). However, the feature extraction, which is run once in the beginning, is still the major bottleneck when it comes to speed. The vertical dotted line indicates the original size of the data set. The vertical dotted line indicates the original size of the data set. Each subset was repeated 3 times, and the speed was averaged across seeds. Error bars represent the standard deviation

Table 1: Detailed description of the extracted features

| Type | Feature | Body part(s) | Description |
|---|---|---|---|
| Intra-animal | Distance | all but ears | *distance in pixels between two body parts of the same animal, e.g., resident snout and resident tail base* |
| | Angular change | all but ears | *angular change within a **bout** of two body parts of the same animal, e.g., snout and tail base* |
| | Speed | all but ears | *speed in pixels/second of an animal measured within a **bout*** |
| Inter-animal | Distance | all but ears | *distance in pixels between two body parts of different animals, e.g., resident snout and intruder snout* |
| | Angular change | all but ears | *angular change within a **bout** of two body parts of different animals, e.g., resident snout and intruder tail base* |

Table 2: Average number of labels during active learning iterations for CalMS21 in Fig. 2b. Averaged across evaluations (20-fold). Total number of available labels (n = 15866); per class: Attack = 1188, Investigation = 12300, Mount = 2378.

| iteration | attack | investigation | mount | total |
|---|---|---|---|---|
| 1 | 91.4 | 165.2 | 97.55 | 354.15 |
| 2 | 112.85 | 316.5 | 127.65 | 557. |
| 3 | 180.35 | 395.65 | 181. | 757. |
| 4 | 230.85 | 470.4 | 255.75 | 957. |
| 5 | 290.55 | 566.7 | 299.75 | 1157. |
| 6 | 353.15 | 646.75 | 357.1 | 1357. |
| 7 | 410.45 | 727. | 400.55 | 1538. |
| 8 | 462.25 | 734.65 | 435.1 | 1632. |
| 9 | 490.75 | 740.85 | 457.4 | 1689. |
| 10 | 509.85 | 770.1 | 481.05 | 1761. |
| 11 | 525.45 | 798. | 504.55 | 1828. |
| 12 | 537.3 | 813.7 | 515. | 1866. |
| 13 | 544.65 | 824.35 | 524. | 1893. |
| 14 | 552.75 | 832.1 | 527.15 | 1912. |
| 15 | 557.9 | 840.9 | 531.2 | 1930. |
| 16 | 561.1 | 849.75 | 532.15 | 1943. |
| 17 | 567.25 | 856.3 | 534.45 | 1958. |
| 18 | 569.35 | 860. | 537.65 | 1967. |
| 19 | 570.5 | 874.6 | 539.9 | 1985. |
| 20 | 576.7 | 875. | 541.3 | 1993. |

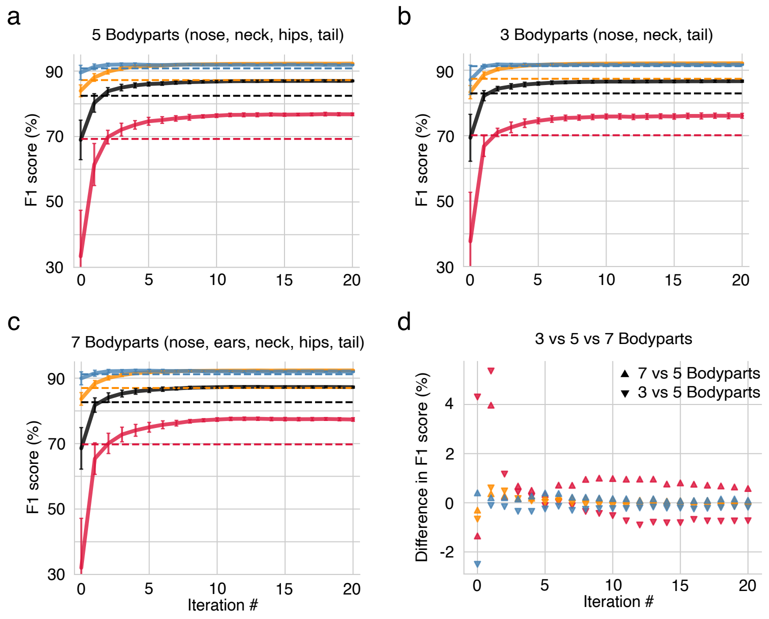


Figure S9: A-SOiD performance done on 3, 5, or 7 keypoints. a) Identical to what was used in the Fig. 2, 5 keypoints were used (nose, neck, two hips, and tail base). b) A-SOiD performance on using just 3 keypoints (nose, neck, and tail base). c) A-SOiD performance on using all 7 keypoints (nose, two ears, neck, two hips, and tail base) d) F1 score difference between 5 keypoints and either all 7 keypoints or a reduced 3 keypoints over active learning iterations. The error bars represent the standard deviation across cross-validations.

Table 3: Average number of labels during active learning iterations for single housed monkey in Fig. 4b. b) per class. Averaged across evaluations (20-fold). Total number of available labels (n = 1181); per class: Climb (C) = 64, Climb (S) = 177, Jump = 50, Rear = 214, Walk = 676.

| iteration | Climb (C) | Climb (S) | Jump | Rear | Walk | total |
|---|---|---|---|---|---|---|
| 1 | 5.9 | 13.85 | 3.05 | 14.55 | 33.65 | 71 |
| 2 | 9.05 | 19 | 4.45 | 18.35 | 35.15 | 86 |
| 3 | 11.7 | 24.1 | 6.45 | 21.45 | 37.3 | 101 |
| 4 | 14.45 | 27.8 | 8.8 | 24.55 | 40.4 | 116 |
| 5 | 15.9 | 31.7 | 11.4 | 28.8 | 43.2 | 131 |
| 6 | 17.5 | 35.55 | 14.35 | 31.95 | 46.65 | 146 |
| 7 | 19.4 | 38.6 | 16.5 | 35.1 | 51.4 | 161 |
| 8 | 20.75 | 41.35 | 19.3 | 39.35 | 55.25 | 176 |
| 9 | 22.2 | 45.45 | 22.15 | 41.6 | 59.6 | 191 |
| 10 | 24 | 48.2 | 25.65 | 44.45 | 63.7 | 206 |
| 11 | 25.35 | 50.55 | 29.15 | 47.7 | 68.25 | 221 |
| 12 | 26.15 | 53.25 | 32.15 | 50.3 | 73.8 | 235.65 |
| 13 | 27 | 55.1 | 35.05 | 52.15 | 79.55 | 248.85 |
| 14 | 27.35 | 57.2 | 36.3 | 54.35 | 83.3 | 258.5 |
| 15 | 27.8 | 58.55 | 37.1 | 55.6 | 85.35 | 264.4 |

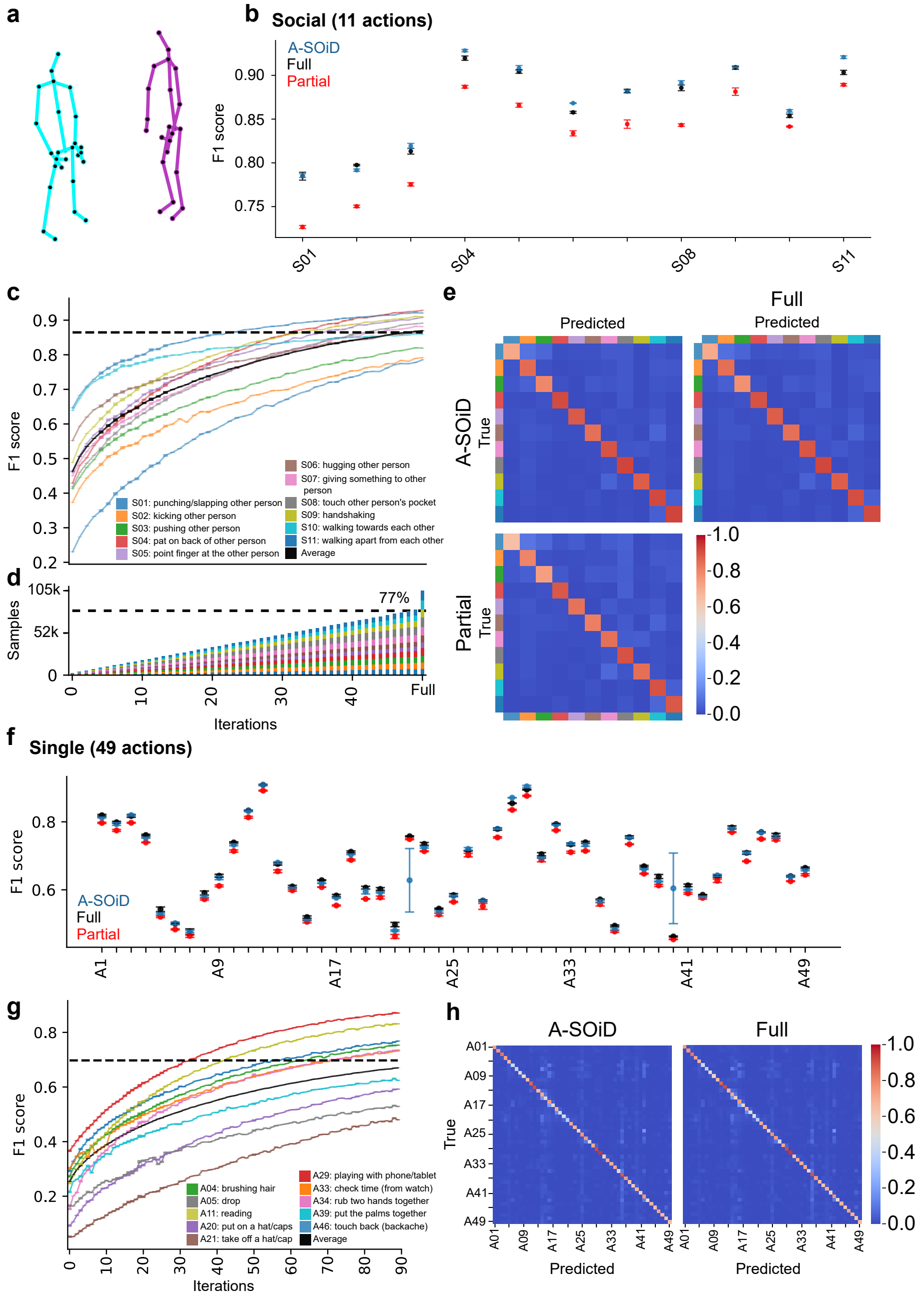


Figure S10: To verify whether our approach works with data sets that contain a higher number of actions, including more complex actions, we selected the public benchmark data set NTU-RGB-D60 and used a subset of the data to evaluate A-SOiD's performance (see Methods). For this, we split the data set into two based on the number of involved subjects. Social human actions (a-e) and Single human actions (g-h). a) Schematic representation of the pose estimation skeletons in the data set. Each skeleton is represented by 25 keypoints.

Figure S10: b) A-SOiD performance (F1 score, 3 cross-validations, blue) on held-out test data of the last iteration (50 iterations; n_samples = 81045, 77% of training set) plotted versus the performance of a classifier trained with the same number of samples (partial, red) and with the full training set (full, black). See Suppl. Table 4 for details. c) A-SOiD performance (F1 score, 3 cross-validations) on held-out test data is plotted against active learning iterations. Black: Unweighted class average. The dashed line demonstrates unweighted class average performance using full training annotations at once. Error bars represent the standard deviation across cross-validations. d) Stacked bar graph of the number of training annotations is plotted against active learning iterations. The right-most bar represents the full annotation count. Bars represent average training samples across 3 seeds. The dashed line indicates the number of samples in the final iteration (77%, n = 81045). e) Confusion matrices where prediction mistakes were being made for the last iteration (A-SOiD, top) and a classifier trained on the same number of samples, randomly sampled from the training set (Partial, bottom). Darker shades along the diagonal indicate better algorithm performance in matching the human rater target. f) A-SOiD performance across all 49 actions (F1 score, 3 cross-validations, blue) on held-out test data of the last iteration (90 iterations; n_samples = 546245, 87% of training set) plotted versus the performance of a classifier trained with the same number of samples (partial, red) and with the full training set (full, black). Error bars represent the standard deviation across cross-validations. See Suppl. Table 5 for details. g) A-SOiD performance (F1 score, 3 cross-validations) on held-out test data is plotted against active learning iterations. Ten representative actions, including the worst and best performing, are shown. Dashed lines demonstrate performance using full training annotations at once. Black: Unweighted class average across all 49 actions. Error bars represent the standard deviation across cross-validations. h) Confusion matrices where prediction mistakes were being made for the last iteration (A-SOiD, left) and a classifier trained on the full training set (Full, right). Darker shades along the diagonal indicate better algorithm performance in matching the human rater target.

Table 4: Performance (f1 score) and standard deviation (SD) of classifiers shown in Supplementary Fig. S10 b.

| | **Full** | | **Partial** | | **A-SOiD** | |
|---|---|---|---|---|---|---|
| **ID** | Performance | SD | Performance | SD | Performance | SD |
| S01 | 0.784755 | 0.004498 | 0.726691 | 0.001744 | 0.785417 | 0.002138 |
| S02 | 0.797479 | 0.000830 | 0.750289 | 0.001442 | 0.791657 | 0.001345 |
| S03 | 0.813283 | 0.003285 | 0.775366 | 0.001919 | 0.819342 | 0.002901 |
| S04 | 0.919655 | 0.002500 | 0.886932 | 0.001605 | 0.928262 | 0.001358 |
| S04 | 0.904438 | 0.002116 | 0.865882 | 0.002340 | 0.908383 | 0.002879 |
| S05 | 0.857722 | 0.001198 | 0.833744 | 0.002959 | 0.868229 | 0.000426 |
| S06 | 0.882259 | 0.001842 | 0.844259 | 0.004797 | 0.881508 | 0.001782 |
| S08 | 0.885776 | 0.003311 | 0.843172 | 0.001480 | 0.891370 | 0.002615 |
| S09 | 0.908822 | 0.001779 | 0.881376 | 0.004265 | 0.909402 | 0.000336 |
| S10 | 0.853744 | 0.001984 | 0.841499 | 0.000650 | 0.859391 | 0.001457 |
| S11 | 0.903328 | 0.002440 | 0.889266 | 0.001410 | 0.920768 | 0.001574 |

Supplementary Movie 1: Video examples of two sub-classes segmented from investigation that reflect anogenital investigation. On the left, one mouse directly approaches the anogenital area of another mouse, irrespective of the incoming angle ("anogenital approach"). On the right, one mouse investigates the anogenital area of another mouse while already being in close proximity to begin with ("anogenital investigation").

Table 5: Performance (f1 score) and standard deviation (SD) of classifiers shown in Supplementary Fig. S10 f.

| | **Full** | | **Partial** | | **A-SOiD** | |
|---|---|---|---|---|---|---|
| **ID** | Performance | SD | Performance | SD | Performance | SD |
| **A1** | 0.819442 | 0.002385 | 0.796839 | 0.001672 | 0.812567 | 0.001161 |
| **A2** | 0.798719 | 0.002707 | 0.774755 | 0.002767 | 0.793360 | 0.003682 |
| **A3** | 0.817240 | 0.003959 | 0.797245 | 0.001202 | 0.819705 | 0.002466 |
| **A4** | 0.761154 | 0.002227 | 0.739470 | 0.000767 | 0.753200 | 0.001238 |
| **A5** | 0.542645 | 0.006779 | 0.520727 | 0.002387 | 0.528742 | 0.002666 |
| **A6** | 0.499930 | 0.001692 | 0.483489 | 0.001611 | 0.501695 | 0.003763 |
| **A7** | 0.476212 | 0.008886 | 0.464406 | 0.004572 | 0.477526 | 0.003236 |
| **A8** | 0.590588 | 0.005973 | 0.572244 | 0.002599 | 0.580375 | 0.002937 |
| **A9** | 0.641428 | 0.004395 | 0.611540 | 0.002809 | 0.634963 | 0.004956 |
| **A10** | 0.739250 | 0.002101 | 0.713968 | 0.003254 | 0.732390 | 0.002744 |
| **A11** | 0.833369 | 0.001089 | 0.813317 | 0.002856 | 0.831550 | 0.001691 |
| **A12** | 0.908902 | 0.001580 | 0.891804 | 0.001415 | 0.908481 | 0.001586 |
| **A13** | 0.676210 | 0.003584 | 0.654570 | 0.004165 | 0.679794 | 0.002706 |
| **A14** | 0.609833 | 0.003098 | 0.598406 | 0.002667 | 0.605849 | 0.000605 |
| **A15** | 0.519747 | 0.003450 | 0.505903 | 0.003233 | 0.511087 | 0.001495 |
| **A16** | 0.628274 | 0.001235 | 0.607823 | 0.000462 | 0.619370 | 0.000712 |
| **A17** | 0.583170 | 0.003209 | 0.553864 | 0.001538 | 0.576420 | 0.001376 |
| **A18** | 0.711927 | 0.002584 | 0.687846 | 0.002438 | 0.704079 | 0.000432 |
| **A19** | 0.606373 | 0.003898 | 0.573175 | 0.000773 | 0.594580 | 0.006857 |
| **A20** | 0.602439 | 0.003943 | 0.577410 | 0.004212 | 0.592391 | 0.001764 |
| **A21** | 0.497774 | 0.006376 | 0.462841 | 0.006023 | 0.480061 | 0.002340 |
| **A22** | 0.757588 | 0.001213 | 0.749169 | 0.001838 | 0.628031 | 0.093389 |
| **A23** | 0.735102 | 0.004099 | 0.713495 | 0.001357 | 0.725084 | 0.002658 |
| **A24** | 0.544558 | 0.001628 | 0.527250 | 0.004348 | 0.534308 | 0.003544 |
| **A25** | 0.584503 | 0.004251 | 0.564411 | 0.001507 | 0.581669 | 0.001513 |
| **A26** | 0.714851 | 0.000811 | 0.702195 | 0.005171 | 0.720227 | 0.003958 |
| **A27** | 0.568215 | 0.002470 | 0.551151 | 0.008724 | 0.567142 | 0.002325 |
| **A28** | 0.778720 | 0.003517 | 0.754152 | 0.002129 | 0.779437 | 0.001900 |
| **A29** | 0.854664 | 0.000877 | 0.835101 | 0.001670 | 0.870936 | 0.000384 |
| **A30** | 0.894811 | 0.001286 | 0.876433 | 0.001731 | 0.904342 | 0.003037 |
| **A31** | 0.704776 | 0.004670 | 0.687973 | 0.005557 | 0.694544 | 0.002008 |
| **A32** | 0.793522 | 0.001907 | 0.774920 | 0.001494 | 0.790992 | 0.000633 |
| **A33** | 0.733493 | 0.003787 | 0.710562 | 0.002925 | 0.735323 | 0.002781 |
| **A34** | 0.739820 | 0.002782 | 0.714350 | 0.001956 | 0.731972 | 0.003885 |
| **A35** | 0.571662 | 0.001403 | 0.557660 | 0.005185 | 0.566687 | 0.004303 |
| **A36** | 0.494327 | 0.002223 | 0.477466 | 0.002772 | 0.485148 | 0.003401 |
| **A37** | 0.755002 | 0.003915 | 0.734060 | 0.001152 | 0.753594 | 0.001703 |
| **A38** | 0.669400 | 0.002429 | 0.647488 | 0.001340 | 0.660641 | 0.000997 |
| **A39** | 0.639115 | 0.006006 | 0.612885 | 0.003208 | 0.624418 | 0.001276 |
| **A40** | 0.462599 | 0.001539 | 0.454338 | 0.000545 | 0.604319 | 0.103844 |
| **A41** | 0.613093 | 0.003835 | 0.589638 | 0.002679 | 0.600291 | 0.001012 |
| **A42** | 0.585293 | 0.000705 | 0.576138 | 0.003395 | 0.579309 | 0.001769 |
| **A43** | 0.639854 | 0.003243 | 0.626623 | 0.005063 | 0.642463 | 0.003316 |
| **A44** | 0.783433 | 0.004746 | 0.769635 | 0.001017 | 0.780849 | 0.002500 |
| **A45** | 0.708916 | 0.004482 | 0.683966 | 0.001086 | 0.708622 | 0.002658 |
| **A46** | 0.769875 | 0.000679 | 0.749415 | 0.002059 | 0.768374 | 0.001420 |
| **A47** | 0.761713 | 0.004371 | 0.747038 | 0.003054 | 0.756043 | 0.003281 |
| **A48** | 0.640362 | 0.002027 | 0.625354 | 0.001046 | 0.639077 | 0.002525 |
| **A49** | 0.664574 | 0.002210 | 0.644277 | 0.002949 | 0.658664 | 0.002684 |
